# Siderophore-mediated iron partition promotes dynamical coexistence between cooperators and cheaters

**DOI:** 10.1101/2022.09.13.507871

**Authors:** Jiqi Shao, Nan Rong, Zhenchao Wu, Shaohua Gu, Beibei Liu, Ning Shen, Zhiyuan Li

## Abstract

Microbes shape their habitats through consuming resources, as well as actively producing and secreting diverse chemicals. These chemicals serve various niche-construction functions and can be considered “public good” for the community. Most microorganisms, for instance, release small molecules known as siderophores to scavenge irons from the extracellular environment. Despite being exploitable by cheaters, biosynthetic genes producing such molecules widely exist in nature, invoking active investigation on the possible mechanisms for producers to survive cheater invasion. In this work, we utilized the chemostat-typed model to demonstrate that the division of the iron by private and public siderophores can promote stable or dynamical coexistence between the cheater and “partial cooperators”, an adaptive strategy with the production of both public and private siderophores. Further, our analysis revealed that when microbes not only consume but also produce resources, this type of “resource partition model” exhibit different stability criteria than that of the classical consumer resource model, allowing more complex systems dynamics.

## Introduction

Microbes interact by shaping their microhabitats (Bajić, Rebolleda-Gómez et al. 2021). Essential to this intricate ecological loop is the “chemical environment”: the substances in the local inhabitants that directly influence and are influenced by microbes(Ley, Peterson et al. 2006, Delmont, Robe et al. 2011). Common examples of such substances include “resources” such as carbon, nitrogen, and oxygen, which are supplied into the extracellular environment and are consumed by microorganisms for their own growth(Boyd and Ellwood 2010, Miransari 2013, Dutkiewicz, Cermeno et al. 2020). In theoretical ecology, the consumer resource model has been utilized for decades to explore the feedback between species and resources(Lafferty, DeLeo et al. 2015), whereby the dimension of the chemical space has been demonstrated to be of key relevance: MacArthur et al. validated the so-called competitive exclusion principle (CEP) that the number of stably coexisting species cannot exceed the number of resources for which they are competing(Hardin and G. 1960, Macarthur and Levins 1964); In the chemical space of dimension two, Tilman et al. demonstrated that the zero-net growth isoclines and the vector of consumption determine the outcome of competition(Tilman 1982); And in a multi-species resource competition system with a minimal chemical dimension of three, Huisman et al. proved that oscillatory and even chaotic dynamics could emerge to permit dynamical coexistence that exceeds the upper bounds of CEP(Huisman and Weissing 1999, Huisman and Weissing 2001). Actually, numerous research efforts have been devoted to conciliating the conflicts between CEP and the apparent biodiversity in nature, most of which incorporate additional factors such as spatial or temporal heterogeneity(Erez, Lopez et al. 2020, Ho, Good et al. 2022). No definitive answer has yet been achieved (Roy and Chattopadhyay 2007, Gupta, Garlaschi et al. 2021).

Microorganisms have their means of lifting the upper limit of competitive exclusion. By leaking metabolic byproducts or actively producing and secreting secondary metabolites, cells can expand the chemical diversity of their habitats(Schmidt, Ulanova et al. 2019). Such activities, in which microorganisms generate new chemical dimensions, have profound implications for microbial ecology (Fischbach and Segre 2016, Estrela, Diaz-Colunga et al. 2022). For example, pervasive cross-feeding through metabolic byproducts has been shown suggested to promote stable coexistence even under a single carbon source(Goldford, Lu et al. 2018). Meanwhile, the vast variety of bacteriocins and antibiotics escalates microbial chemical warfare (Czaran, Hoekstra et al. 2002, Granato, Meiller-Legrand et al. 2019, Niehus, Oliveira et al. 2021). For microbial cooperation, various quorum sensing molecules coordinate the collective decision-making (Ross-Gillespie and Kümmerli 2014), and niche-construction molecules such as siderophores or secretory proteases serve as “public goods” that improve the microenvironment for the whole population(Leventhal, Ackermann et al. 2019, Smith and Schuster 2019). In ecological theories, however, the production of secondary metabolites confronts the same dilemma as public goods in “the tragedy of the commons” (Hardin 1968): if a cheater strain can always take advantage of public goods without paying for them, how can the cooperator strains accountable for these extra chemical dimensions remain competitive?

Big questions in biology may find diverse solutions in particular systems. Siderophore, a diverse family of microbial secondary metabolites for iron-scavenging, serves as a nice model system to study the self-generated chemical dimensions and the game between cooperators and cheaters (Cremer, Melbinger et al. 2019). Iron is among one of the most limiting resources for microbes(Andrews, Robinson et al. 2003). It is necessary for processes like energy metabolism and DNA synthesis(Andrews, Robinson et al. 2003), yet the concentration of bioavailable iron is orders of magnitude lower than what is required for normal microbial growth in most environments (Boyd and Ellwood 2010, Emerson, Roden et al. 2012). Most microorganisms acquire iron via siderophores, a type of small molecule with a strong iron-binding affinity (Buckling, Harrison et al. 2010). Previous research suggested that siderophores are secreted into the extracellular environment to chelate trivalent iron, and that the iron-siderophore complex is then acquired by membrane receptors so that cells can absorb iron (Kramer, Özkaya et al. 2020). The production of siderophores, on the other hand, comes at a considerable metabolic burden and slows down microbial growth (Lv, Hung et al. 2014, Sexton and Schuster 2017). Therefore, siderophore acts as a costly public good that invokes intricate ecological games. Many theories have been proposed to explain the prevalence of siderophore synthesis pathways (Butaitė, Baumgartner et al. 2017, Gu, Wei et al. 2020), most of which entail spatial factors that facilitate kin selection or group selection (Allison 2005, Ross-Gillespie, Gardner et al. 2007, Julou, Mora et al. 2013). Yet, in well-mixing environments such as the ocean, microorganisms continue to actively produce siderophores of various types (Cordero, Ventouras et al. 2012, Hagstrom and Levin 2017).

Recent experiments indicated that private siderophores, which are solely accessible to their cooperators, may be crucial to the survival of siderophore-producing microorganisms (Scholz and Greenberg 2015). In some organisms, during the multi-step process of producing siderophores, certain modifications can transform the secretory molecules into a membrane-attached form (Martinez, Carter-Franklin et al. 2003, Scholz and Greenberg 2015, Niehus, Picot et al. 2017). Such privatization avoids diffusion losses and cheater exploration, resembling the “snowdrift” scenario in game theory (Souza, Pacheco et al. 2009): Despite that cheaters continue to take advantage of public goods without paying, cooperators have prioritized access to the goods to preserve coexistence (Gore, Youk et al. 2009). Nevertheless, membrane-attached siderophores suffer a considerably lower diffusion radius in scavenging iron (Leventhal, Ackermann et al. 2019, Kramer, Özkaya et al. 2020). Whether the marginal benefits conferred by membrane-attached siderophores are adequate to provide cooperators with a sufficient advantage over cheaters has not been quantitatively assessed by ecological models. Many questions remain to be systematically explored by mathematical formulation, such as whether and how cooperators and cheaters coexist, how different allocation strategies between membrane-attached and public-shareable secretory siderophores change the system dynamics, and which strategies might be “optimal” considering within species and between species. Further, may this siderophore-mediated iron competition helps address a deeper question in ecology, namely whether such self-generated resource dimensions change the properties of the system fundamentally?

In this work, we utilized the chemostat-type “resource partition model” to examine the siderophore-mediated iron competition, taking into account the allocation of limited cellular resources between the biomass accumulation and the production of membrane-attached (private) and public-sharable (public) siderophores. With the inclusion of private siderophores into the resource allocation strategies, new classes of strategies emerge, such as “partial cooperators” that produce both types of siderophores, and “self-seekers” that produce only membrane-attached siderophores. We confirmed mathematically that private siderophores play a vital role in coexistence: private siderophores provide partial cooperators with an advantageous growth zone over cheaters to enable coexistence, in contrast to pure cooperators who always lose to cheater invasion. Interestingly, such two-species coexistence can occur via dynamical oscillation. Further stability analysis revealed that the action of species creating new resource dimensions, i.e. the production of siderophores in our model, modifies the stability criteria of the classical consumer resource model, allowing for rich dynamics in the parameter regions of stable equilibrium for the classical model. In summary, our model in iron competition suggested that the division of the iron resource by siderophores raises the upper-bounds of coexistence, and the privatization of siderophores in cellular strategies provides advantages to partial cooperators to realize coexistence. Our analysis of the stability criteria revealed that the microbial niche-construction by creating new resource dimensions adds more dynamics than the classical model, which may contribute to the diversity and complexity of the microbial world.

## Result

### A resource partition model with trade-offs between growth and siderophore-production

In the field of theoretical ecology, the consumer resource model mimics ecosystems with constant nutrient supplies and extensive mixings, such as lakes and oceans, where species compete by consuming the supplied resources. Nonetheless, in the microbial world, species not only consume but also produce resources that can be shared by the community, such as siderophores for iron scavenging. To examine the ecological consequences of siderophore production and privatization, we developed a mathematical model resembling a chemostat with trade-offs between growth and siderophore production, with the following three assumptions:

1. Chemostat-typed resource partition: we used a chemostat-type model, in which the volume is maintained constant by maintaining the same rate of in-and-out fluxes. Parameters about the concentration of iron in the influx *R*_iron,supply_, together with the dilution rate *d*, are referred to as “chemostat conditions” (Fig. 1A). Within the chemostat, the chemical environment cells directly facing can be quantified by two variables: concentration of the iron (*R*_iron_) and of the public siderophores (*R*_sid_). They serve as both “resources” for microbial growth; meanwhile, the public siderophore is also a “product” released by bacteria. We coined the term “resource partition model” to refer to models in which organisms generate more chemical dimensions than are supplied from the external influx, and partition the externally supplied resources by these secondary metabolites.
2. The growth rate linearly scales with the iron uptake rate: Assuming iron was the limiting resource, we set the growth rate to a value that linearly scales the total iron fluxes obtained via two types of siderophores: the flux from secretive public siderophores (*J*_public_, depicted on the left side of Fig. 1B) and the flux from membrane-bounded private siderophore (*J*_private_, depicted on the right side of Fig. 1B).
3. Trade-offs in resource allocation between growth and siderophore-production: Given the limited amount of proteins and energies in a microorganism, we assumed that there are trade-offs between different biological processes. In an iron-limited environment, we focused on resource partitioning into biomass accumulation and production of private and public siderophores. Each partition corresponds to an allocation strategy 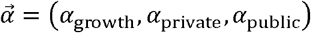, where *α*_private_ and *α*_public_ denote the percentage of resources used to produce private and public siderophores, respectively, and *α*_growth_ donates the percentage of resources devoted to biomass accumulation. The summation of all allocations was fixed as *α*_σ,growth_ + *α*_σ,private_ + *α*_σ,public_ = 1. Various species may differ in their abilities to produce and scavenge siderophores, but different strains σ only differ in their resource allocation strategies 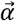. Under this trade-off, all possible strategies of a species can be located in a ternary graph (Fig. 1C). There are four types of typical strategies (Fig. 1C-D): pure cooperators, the pure cheater, partial cooperators, and self-seekers. Pure cooperators produce public siderophores but not private siderophores (*α*_private_ = 0, *α*_public_ > 0), while the pure cheater allocates all resources to growth and exploits only the public siderophores (*α*_growth_ = 1). On the other hand, partial cooperators (*α*_public_ > 0, *α*_private_ > 0) and self-seekers(*α*_public_ = 0, *α*_private_ > 0) have access to both public and private siderophores due to non-zero *α*_private_. The difference between the two strategies is that partial cooperators still produce public siderophores while self-seekers do not. In the ternary graph (Fig. 1C), pure cooperators’ strategies span the triangle’s base, while the pure cheater’s strategy locates on the right-vertex. The right side is occupied by self-seekers, and each strategy contained within the triangle can be considered a partial cooperator.

**Figure 1.**
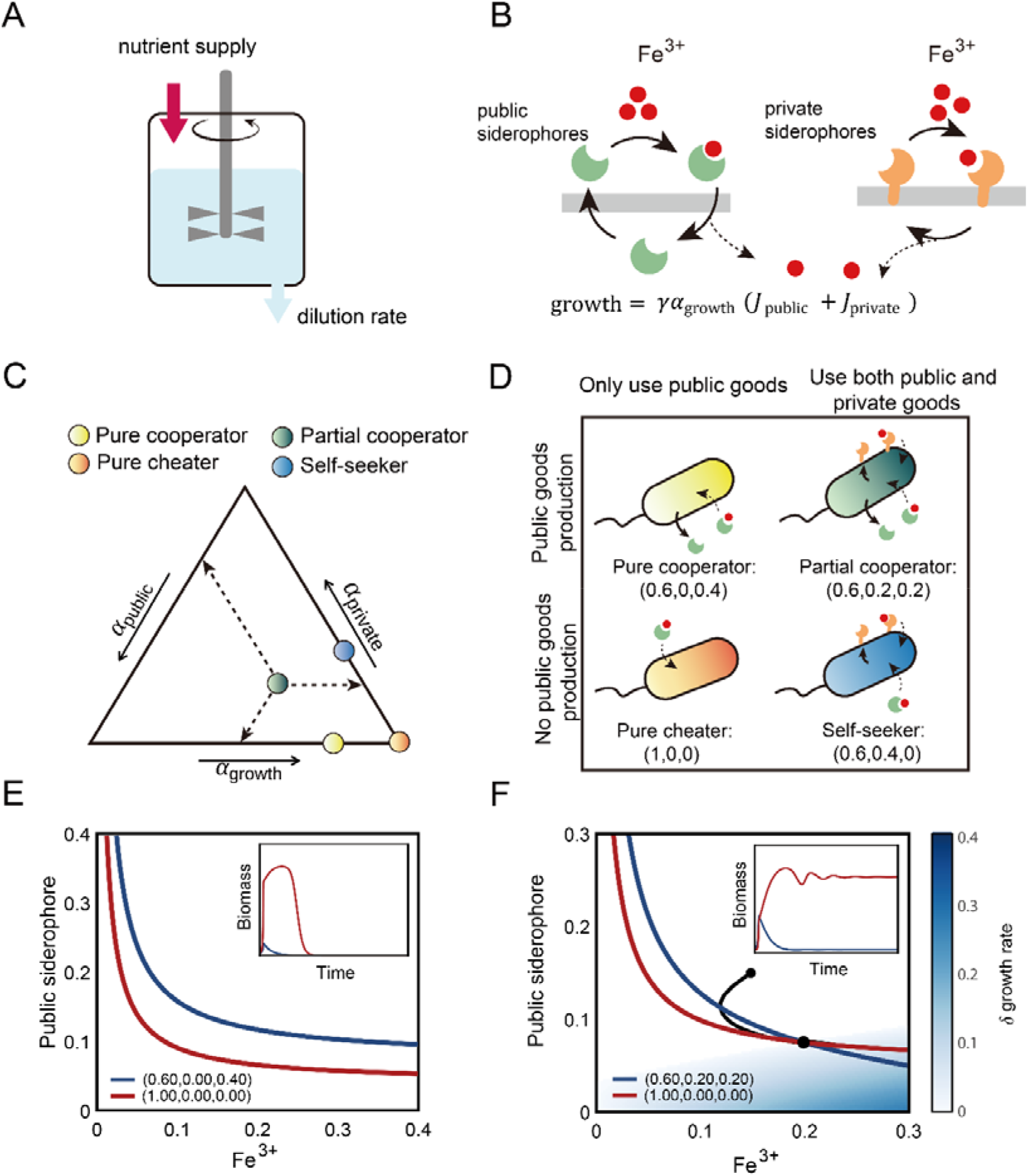
Privatization of siderophore enables the coexistence between partial cooperators and pure cheaters. (A) The schematic diagram of a chemostat model. (B) The schematic diagram showing two iron uptake fluxes from the public siderophores (left) and private siderophores (right). (C) The strategy space (the strategic phase diagram) by ternary plot, showing all resource allocation strategies 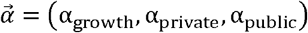. The model has four typical strategies, distinguished according to whether to produce public siderophores and whether to use private siderophores. Pure cooperators (yellow) only produce and use public siderophores, while the pure cheater (orange) only use public siderophores and put all resources into α_growth_; partial cooperators (green) produce and use both public and private siderophores, while self-seekers (blue) only produce private siderophores and use both public and private siderophores. (D) The schematic illustration of the four typical strategies shown in (C). (E-F) The growth contours in the chemical space, with the concentration of Fe^3+^ by the x-axis and the concentration of public siderophore by the y-axis. (E) shows the growth contours of a pure cooperator and the pure cheater, and (F) shows the growth contours of a partial cooperator and the pure cheater (color scheme same as that in (C) and (D)). The blue area represents a growth-advantageous zone in which the partial cooperator outperforms the pure cheater. Inserts are the biomass time-courses of species competing in the chemostat.

With the three assumptions above, the microbe σ with biomass concentration *m*_σ_ and strategy 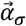 shapes its chemical environment in two ways:

1. Producing public siderophore with the out flux 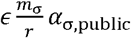, where *ϵ* is the production coefficient of public siderophores, and the constant *r* represents the biomass per cell volume (Details in SI). Assuming the public siderophores can be fully recycled for simplicity, the changing rate for *R*_sid_ can be written as:

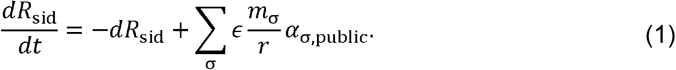
2. Consuming iron by public siderophores and private siderophores. For iron-uptake, the fluxes of *J*_σ,private_ and *J*_σ,public_ take the Monod form with the environmental iron concentration *R*_iron_. We also assumed that both fluxes linearly scale with the concentration of corresponding siderophores. Taken together, there are:

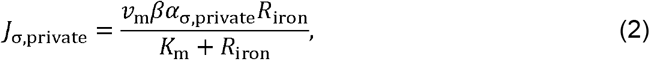

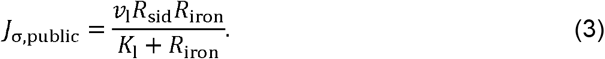 In the equations above, *v*_m_, and *v*_l_ are the rate coefficients for the two fluxes; *K*_m_ and *K*_l_ are the rate coefficients for the two fluxes; *K*_m_ and *K*_l_ are the affinity coefficients of the two kinds of siderophores for intaking iron; *β* is the efficiency coefficient of the private siderophores. In total, the changing rate for iron is written as:

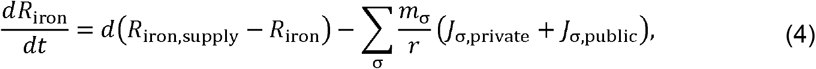 It can be proved that other forms of iron uptake-fluxes do not qualitatively affect the result of this work (See Supplementary Material, Section 4.2).

Meanwhile, the microbe σ has its growth rate affected by its chemical environment [*R*_iron_, *R_sid_*] and allocation strategy 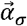 as:

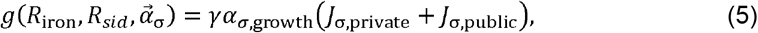

where *γ* here represents the growth coefficient.

In this iron-partition model, the changing rate for the biomass for microbe σ can be written as:

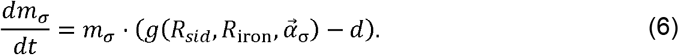

### Privatization of siderophore enables the coexistence between partial cooperators and pure cheaters

The graphical approaches developed for consumer resource models enabled an intuitive assessment of ecological consequences in the chemical space (Tilman 1982) (Fig. S1). This approach consists of two elements: the growth contour, i.e. the zero net growth isoclines (ZNGI), and the consumption vector. After setting Eq. (6) to zero for a single subpopulation, we obtained the growth contours. The growth contour shows all possible [*R*_iron_, *R*_sid_] that the microbe could reach the steady-state growth as the dilution rate. Any environment above the growth contour belongs to the “invasible zone” where the microbe can invade (Li, Liu et al. 2020).

This graphical approach demonstrates intuitively why a pure cooperator cannot coexist with a pure cheater. For two strains, the possible chemical environment allowing for stable coexistence must locate at the intersection of their growth contours, indicated by the “*”. As illustrated in Figure 1E, the growth contour of the pure cheater entirely encloses the growth contour of the pure cooperators without an intersection. Intuitively, it is due to that the pure cheater grows faster than the pure cooperator in any chemical environment: they have the same iron-influx by public siderophores *J*_public_, yet the cooperator invests less in *α*_growth_ (See rigorous proof in the Supplementary Material, Section 2.2). However, because the pure cheater can only utilize public siderophores and cannot survive independently, it can be observed in the simulation that the pure cheater eventually takes the entire population and becomes extinct due to a lack of public siderophores. This approach resembles “the tragedy of the commons”.

Investing in private siderophores can change the game. The transition from pure cooperators to partial cooperators enables the growth contours to intersect with those of pure cheaters, hence enabling coexistence (Fig. 1F). In regions where the public siderophore is abundant but iron is scarce (upper-left region of Figure 1F filled in white), the cheater still retains a growth advantage over the partial cooperator; however, in regions where the public siderophore is scarce but free iron is abundant (lower-right region of Figure 1F filled in blue-green color), the partial cooperator gains a growth advantage over the pure cheater because their membrane siderophores sustain iron-dependent growth. Under this model setting, the game enters the regime of “snowdrift” (Souza, Pacheco et al. 2009): despite the fact that producing siderophores incurs costs, cooperators gain by holding portions of them as private goods, particularly in environments where public siderophores are scarce. The fact that the superior strategy varies according to the chemical environment enables the intersection of their growth contours, which is necessary for coexistence.

Mathematically, it can be analytically proven that the presence of intersection between growth contours of strain 1 and strain 2, under the setting that strain 1 invests less in its own growth *α*_1,growth_ < *α*_2,growth_, requires:

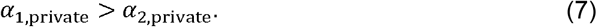

Equation (7) shows that for any two strains to coexist, the strain that invests fewer resources in growth must invest more in private siderophores (Details can be found in Supplementary Material, Section 2.3).

Meanwhile, the non-zero biomass of two strains imposes constraints on the iron supply concentration *R*_iron,supply_ in that

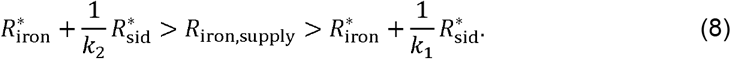

with 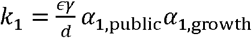, 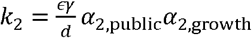.

Graphically, inequality Eq. (8) requires the supply point [*R*_iron,supply_, 0], locating within the region bounded by the reverse extensions of the consumption vectors in the chemical space (See Supplementary Material, Section 2 for detailed proofs)

However, the presence of the intersection of growth contours is only one of the necessary conditions for coexistence; the feasibility of coexistence is further determined by how each strategy shapes the chemical environment for the whole community (Li, Liu et al. 2020). In the system illustrated in Figure 1F, where the partial cooperator coexists stably with the pure cheater (competition dynamics shown in insert), the partial cooperator creates a public-siderophore-rich environment in the upper-left area of the chemical space where the cheater can invade. Meanwhile, the pure cheater cannot survive on its own and has no corresponding stable point, but its competition for iron reduces the abundance of the partial cooperator, causing an environment deficient in public siderophores to favor the partial cooperator. This interaction between the partial cooperator and the pure cheater resembles mutual invasion and allows for stable coexistence.

Such stable coexistence does, however, occur under a specific chemostat condition (quantified as the iron influx *R*_iron,supply_, and the dilution rate *d*). The presence of an area in the chemical space in which the partial cooperator thrives prompted us to wonder whether other chemostat conditions would allow for even more interesting ecological dynamics.

### The partial cooperators and cheaters can generate rich ecological dynamics

In classical consumer resource models, oscillatory dynamics can be generated among a minimum of three species that preferentially consume the resources for which they have intermediate requirements (Huisman and Weissing 1999, Huisman and Weissing 2001). To our surprise, oscillations between the partial cooperator and the pure cheater can be detected in this resource partition model at intermediate levels of iron supply (Fig. 2A). In comparison to the steady coexistence depicted in Figure 1F, the partial cooperator generates an environment susceptible to cheater invasion. The intersection of the two growth contours, on the other hand, becomes unstable.

**Figure 2.**
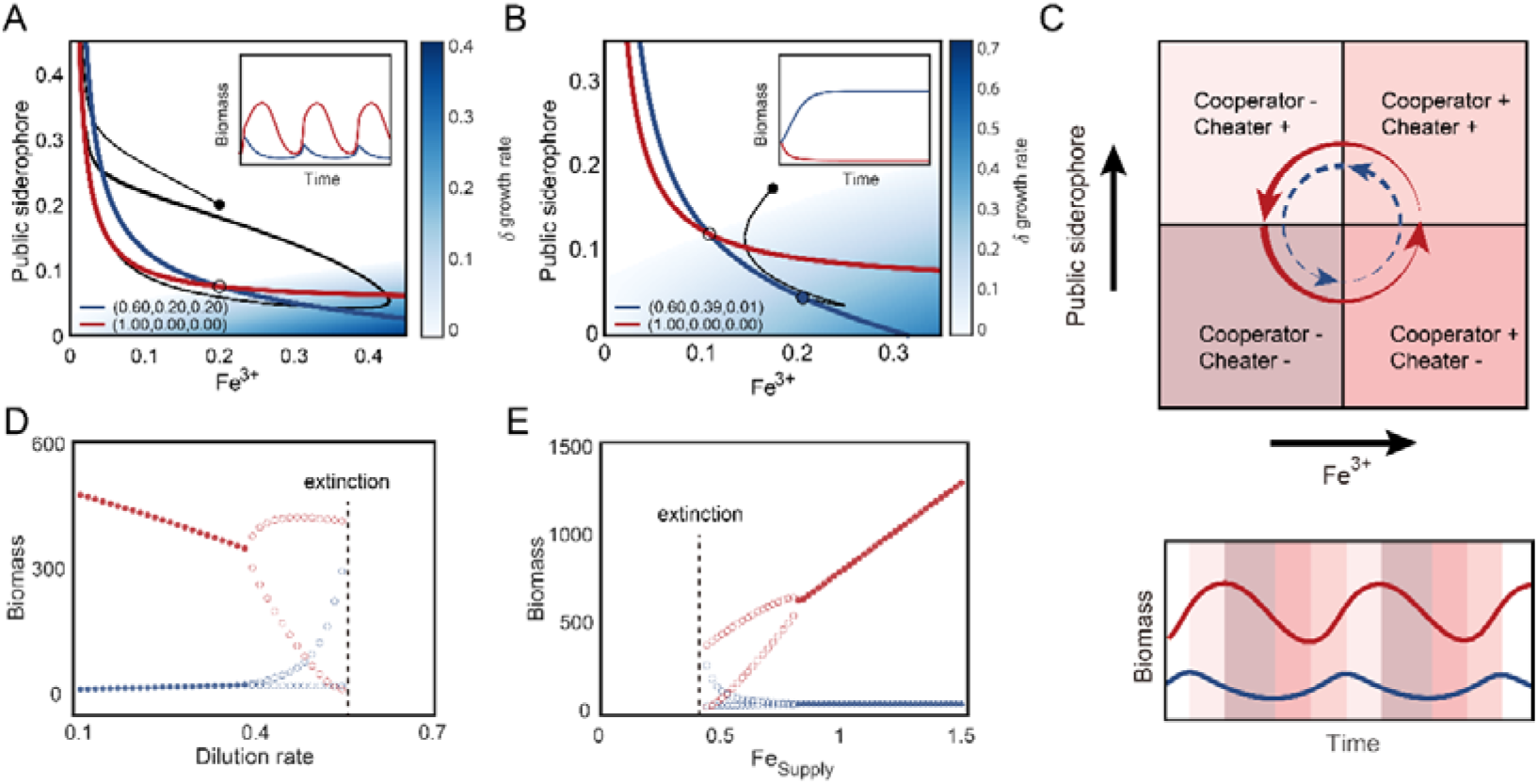
The partial cooperators and cheaters generate rich ecological dynamics. (A) The partial cooperators can oscillate with the cheater. In the chemical space, the intersection between the growth contours of a partial cooperator (blue) and a pure cheater (red) is represented by a black circle, and the dynamical trajectory beginning is represented by a black line. The background color indicates the difference in growth rate between partial cooperators and cheaters. Insert shows the competition dynamics of the two species. (B) The partial cooperators can exclude the cheater. Same as (A), but the partial cooperators allocate more resources to private siderophores. (C) The phase diagram illustrates the mechanism of oscillation shown in (A). In the upper panel, the chemical space is divided into four regions with varying relative finesses (growth rate relative to dilution rate) between the cheater and the partial cooperators, denoted by the symbols + and −. The background color of the lower panel corresponds to the regions in the chemical space depicted in the upper panel. (D-E) Bifurcation diagrams for the system in (A) as the dilution rate (D) and the Fe^3+^ supply changes (E). Empty circles represent the minimum and maximum of the oscillation, while solid dots represent a steady state.

Figure 2C explains the oscillation visually with a phase diagram. In the diagram’s upper-left quadrant, where the public siderophore concentration is high and the iron concentration is low, the rapid proliferation of cheaters effectively eliminates partial cooperators (cooperator −, cheater+). The decline in partial cooperators results in a decrease of public siderophores, which forces the system into the lower-left quadrant, where the abundance of both partial cooperators and cheaters falls (cooperator−, cheater−). Reduced total population promotes iron concentration recovery, allowing the system to enter the advantageous growth zone of partial cooperators, where public siderophores remain low but iron is abundant. Iron influx through private siderophores causes partial cooperators to resume positive growth in this lower-right quadrant, whereas pure cheaters continue to fall (cooperator+, cheater−). The rise of partial cooperators increases the concentration of public siderophores, hence inhibiting the decrease of pure cheaters and restoring their growth, as shown in the upper-right quadrant (cooperator+, cheater+). However, the continued growth of pure cheaters depletes iron, forcing partial cooperators into negative net growth again (cooperator−, cheater+). The preceding process generates oscillations, mostly owing to the private siderophores providing an advantageous growth zone for partial cooperators in the absence of public siderophores.

When the partial cooperator is more capable of establishing a high-iron, low-public-siderophores environment, the dynamics become even more skewed in favor of partial cooperators. For instance, if the steady-state environment formed by partial cooperators goes outside the development contour of pure cheaters, the partial cooperators can effectively exclude the pure cheater (Fig. 2B).

Scanning across the chemostat conditions revealed the bifurcation into and out of oscillation. In the system with one pure cheater and one partial cooperator, as the dilution rate increases, a Hopf bifurcation drives the transition from stable coexistence into oscillation (Fig. 2D-E). Similarly, by increasing the iron supply concentration, the oscillation shifted into stable coexistence. The phase diagram of chemostat conditions (Fig. S2) demonstrates that a “better” environment (low dilution rate or high iron supply) reduces the relative difference in growth between partial cooperators and cheaters, thereby increasing the tendency of coexistence; on the other hand, a harsher environment (high dilution rate or low iron supply) increases the relative difference in growth between the two, thereby increasing the likelihood of oscillations.

Oscillatory dynamics demand a minimum of three species in a classical consumer resource model (Huisman and Weissing 1999, Huisman and Weissing 2001). The observed oscillation between the pure cheater and the partial cooperator not only sheds light on the stability of cooperation in an iron-related snowdrift game, but also motivates us to dig deeper into the effect of the self-generated chemical dimension on the resource partition models.

### The self-generated chemical dimension changes the stability criteria of classical consumer resource models

Similar to a classical consumer resource model (represented by Tilman’s model (Tilman 1982)), the generalized form of microbes interacting with two resources *R*_1_ and *R*_2_ can be represented as:

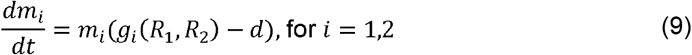

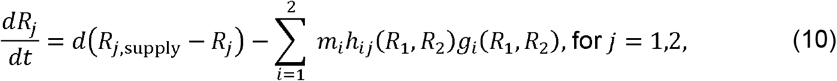

where *R*_*j*,supply_ is the supply concentration of resource *j*, and *h_ij_* is the function describing the amount of resource *j* impacted by microbe *i* per-biomass.

When *h_ij_* describes resource uptake and has a positive sign. Eq. (9)–(10) above represent the classical consumer resource model. When at least one of the *h_ij_* describes resource production and exhibits a negative sign, these equations represent the broader “resource partition model” and exhibit different criteria for the stability of coexistence.

The stability of the fixed point (*) can be deduced from the Routh-Hurwitz (RH) criterion (May and Allen 1977). From the Jacobian matrix of the model:

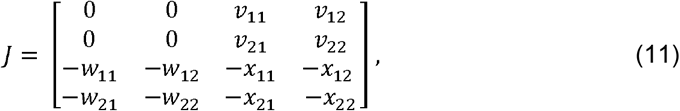

with

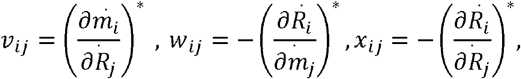

For simplicity, we set the abbreviation as:

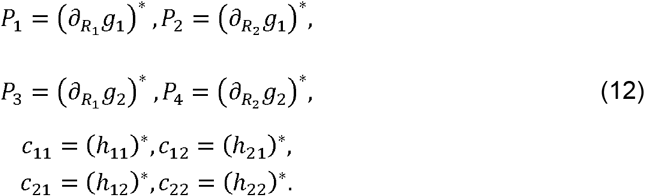

Here, *P* can be interpreted as the growth rate dependency on “resources” at the fixed point, and *c_ij_* represents how microbe *i* impacts resource *j* at the fixed point.

So we have elements in the Jacobian matrix expressed as:

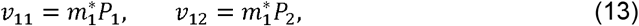

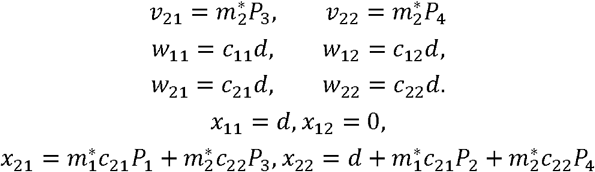

At the steady state, 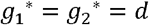. With definitions in Eq.(12)–(13), the characteristic equation of eigenvalue λ for the Jacobian matrix can be expressed as:

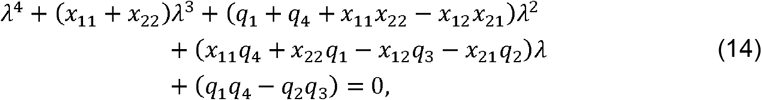

the coefficient of the quartic equation are *a*_0_ = 1, *a*_1_ = *x*_11_ + *x*_22_, *a*_2_ = *q*_1_ + *q*_4_ − *x*_12_*x*_21_ + *x*_11_*x*_22_, *a*_3_ = *q*_4_*x*_11_ − *q*_3_*x*_12_ − *q*_2_*x*_21_ + *q*_1_*x*_22_, *a*_4_ = −*q*_2_*q*_3_ = *q*_1_*q*_4_, with *q_i_* defined as:

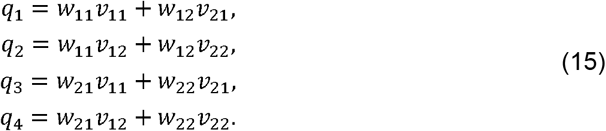

The Routh-Hurwitz (RH) criterion states the necessary and sufficient condition for the stability of a dynamical system (May and Allen 1977): for a quartic equation for the eigenvalue λ, *a*_0_λ^4^ + *a*_1_λ^3^ + *a*_2_λ^2^ + *a*_3_λ + *a*_4_ = 0, if the fixed point for coexistence is stable, then the coefficients must satisfy: (1) *a*_1_ > 0, (2) *a*_4_ > 0, (3) *a*_3_ > 0, (4) 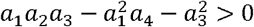.

Based on the four RH criteria, we compared the difference between the classical consumer resource model and the resource partition model on its stability. Details on the comparison can be found in Supplementary Material Section 3.1.

First, the criteria (1) *a*_1_ > 0 holds for both the classical consumer resource model and the iron-partition model.

Second, The criteria (2) *a*_4_ > 0 is central to the classical model. By definitions in Eq. (12), the original form *a*_4_ = −*q*_2_*q*_3_ + *q*_1_*q*_4_ expamds into:

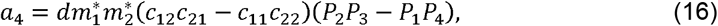

There are two possible situations for (*c*_12_*c*_21_ − *c*_11_*c*_22_)(*P*_2_*P*_3_ − *P*_2_*P*_4_) > 0: situation 1:

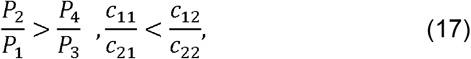

or situation 2:

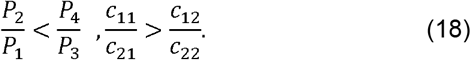

In the classical consumer resource model, these two inequalities can be interpreted as “each species must consume more of the one resource which more limits its own growth rate.”

In the resource partition model, using the iron-partition model at Eq. (1)–(6) as the example, parameters in Eq.(12) can be specified into:

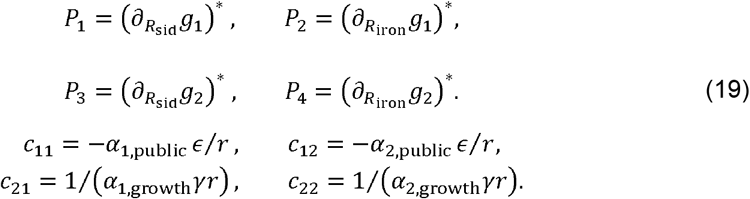

Then it can be proven that under the setting of *α*_1, growth_ < *α*_2, growth_, the criteria (2) *a*_4_ > 0 requires that:

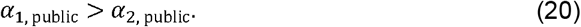

In the iron-partition model, Eq. (7) and Eq. (20) suggest that, if two strains should stably coexist, not only the strain that invests fewer resources in growth must invest more in private siderophores, but also play a more cooperative role.

The remaining two criteria, (3) *a*_3_ > 0 and (4) 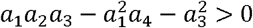, differentiate between the classical model and the iron-partition model. By definitions in Eq. (12)–(15), equations for these two criteria are the additive or multiplicative combination of *a*_4_ with other positive definite terms *c*_11_ and *c*_12_ (details in Supplementary Material, Section 3):

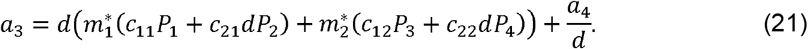

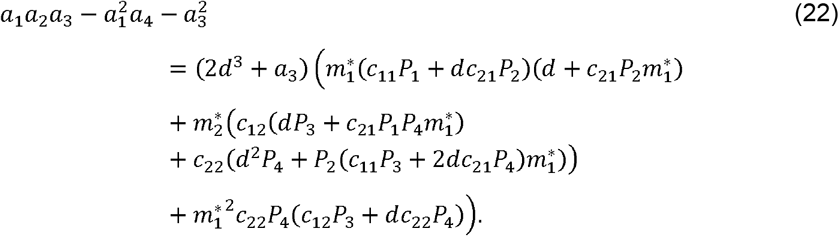

In the classical consumer resource model, *c_ij_* describes the consumption of species *i* on resource *j*, which is always positive. Consequently, once the criteria (2) *a*_4_ > 0 holds, criteria (3) *a*_3_ > 0 and criteria (4) 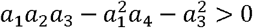 are satisfied automatically. Actually, the determination of the fixed point stability only depends on the tangent slope of growth contours 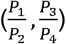 and the slope of the consumption vector 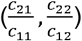. If the criteria in Eq.17–18 are satisfied, the coexistence is stable as long as the supply vector locates within the sector area formed by the reverse extension of the two consumption vectors.

However, in the iron-partition model, the fulfillment of criteria (2) no longer guarantees the satisfaction of criteria (3) and (4). Even when Eq.(17)–(18) are satisfied and the supply vector locates within the sector formed by consumption vectors, changes in the iron-supply can still change the stability of the growth contour intersection, driving the system into other forms of dynamics instead of stable coexistence. As shown in Figure 3A, in the chemical space, the growth contours of a partial cooperator and the pure cheater intersect. At the crossing point, the consumption vector of the partial cooperators points to the upper-left direction, suggesting it consumes iron while creating public siderophores.

**Figure 3.**
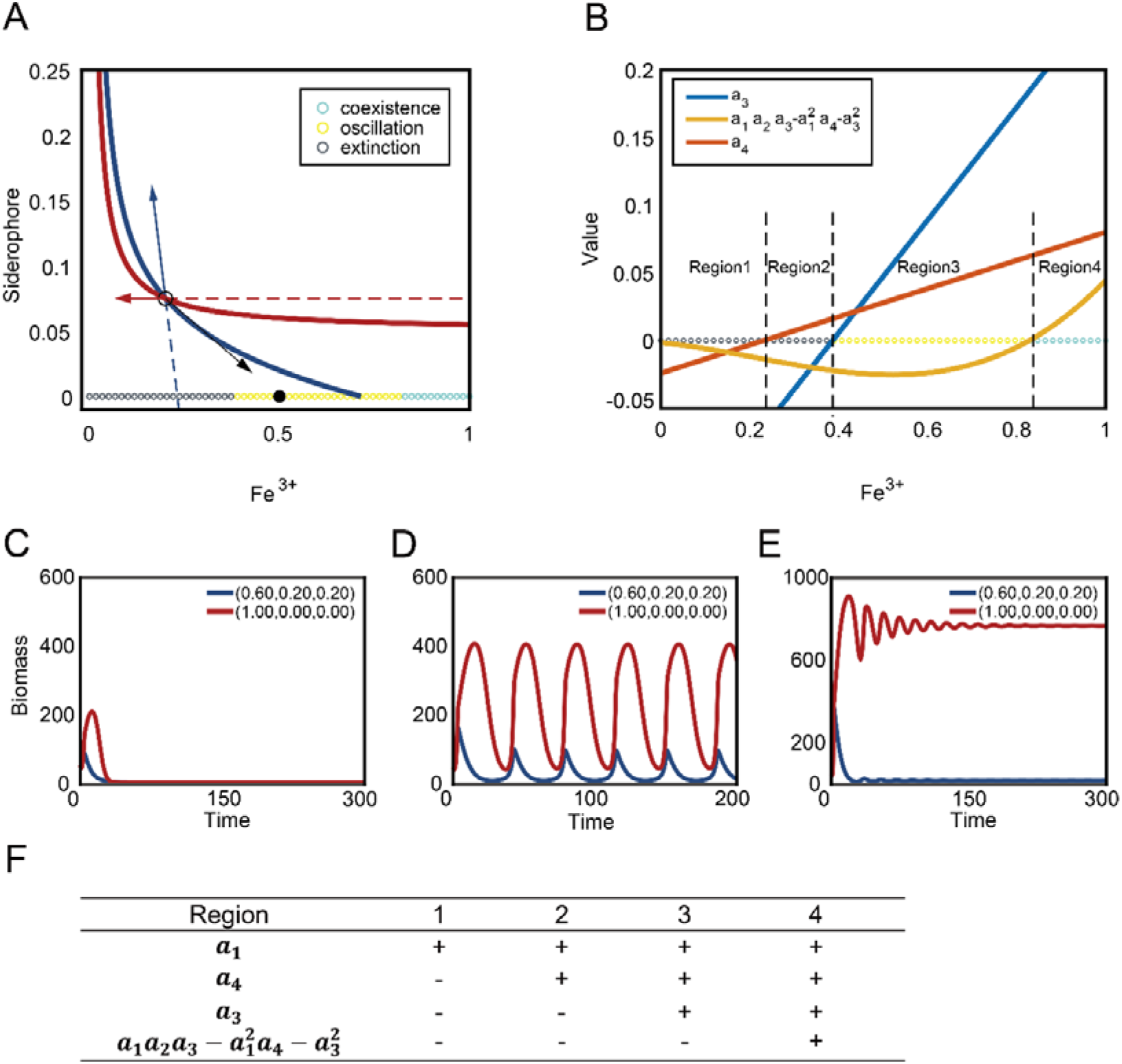
The self-created dimension enables oscillation and coexistence. (A) The growth contours of a partial cooperator (blue) and the cheater (red), and their consumption vectors at the intersection. Different types of dynamics induced by different supplies of Fe^3+^ are indicated by the colors of circles along the x-axis. Black indicates the extinction of both species; Yellow indicates oscillatory dynamics; Blue indicates steady-state coexistence. (B) The values of three stability criteria as the iron supply increases along the x-axis. The signs of three stability criteria divide the x-axis into four regions, as indicated by the text. (C-E) Exemplary time-courses of the three types of dynamics induced by different supplies of Fe^3+^, such as extinction (C), oscillation (D), and coexistence (E). (F) The signs of all four stability criteria in four regions shown in (B).

The consumption vector of the pure cheater horizontally points to the left, as it solely consumes iron but does not contribute to public siderophore. These sectors formed by two consumption vectors overlap a region on the iron-supply axis (x-axis). Nevertheless, within this region of iron-supply, the stability of the fixed point changes: when the iron supply is low, the system will still collapse to extinction even if the supply (black circles, Fig. 3C) locates within the sector zone; oscillation begins as the iron supply increases (yellow circles, Fig. 3D); and eventually, stable coexistence can be reached when the iron supply is high (blue circles, Fig. 3E).

The collapse of the system in Figure 3C can be attributed to the changes in criteria (3) *a*_3_ > 0. Due to the negative signs of *c*_11_ and *c*_12_, the first term of *a*_3_, 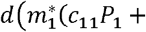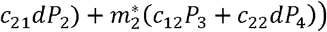 now has an uncertain sign. If 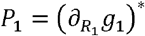 or 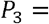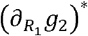, are sufficiently large, indicating that strains are highly sensitive to free iron, the first term might be negative, and is able to drive the whole equation of *a*_3_ into the negative regime. Moreover, the sign of the first term also relies on 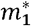 and 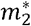, both of which are dependent on the iron-supply concentration. As shown in Region 2 in Figure 3B, the sign of *a*_3_ remains negative when *a*_4_ is positive, which contributes to the extinction of both species even when the iron supply falls within the sector formed by the reverse extension of consumption vectors (Fig. 3C).

Similarly (See Supplementary Materials, Section 3.2), with negative *c*_11_ and *c*_12_, criteria (4) in the iron-partition model now has an uncertain sign, with complex dependance on 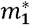 and 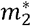. When criteria (4) remains negative (Region 3) while the other criteria are fulfilled(Region 3, Fig. 3B), the fixed point remains unstable, and the system oscillates.

In the classical model, due to the positive sign of *c*_11_ and *c*_12_, Region 2 and Region 3 in Figure 3F would not exist. As the existence of Region 2 and Region 3 is the result of negative *c*_11_ and *c*_12_, such extinction and oscillation dynamics are not exceptional. Two partial cooperators with distinct allocation strategies may also undergo similar transitions, with the dynamics experiencing extinction, oscillation, coexistence, and exclusion as the iron supply increases (Fig. S3).

### Comprehensive assessment of strategies under within-species and between-species competitions

In our model, distinct strains within the same species are distinguished by distinct resource allocation strategies, and their public siderophores can be shared. Parameters such as the cost of siderophores and the growth coefficient can vary between species. In addition, we assumed that siderophores produced by different species can not be shared. We systematically evaluated the competitiveness of strategies within and between species based on these assumptions.

First, the competition between all potential strategies against the pure cheater of the same species exhibits complex dynamics (as illustrated in the ternary plot of Fig. 4A): the pure cheater does trigger extinction for highly cooperative strains (high in *α*_public_ and low in *α*_growth_). However, over a broad range of strategy space, partial cooperator strategies can coexist stably or dynamically with pure cheater strategies. Moreover, this coexistence can exist in a wide range of different parameters and public siderophore production and recycling sets (Fig. S5, S6). Self-seeker strategies can even result in the exclusion of the pure cheater. In terms of coexistence, increasing investment in public siderophores increases the likelihood of oscillations with the cheater, whereas increasing investment in private siderophores increases the tendency of stable coexistence.

**Figure 4.**
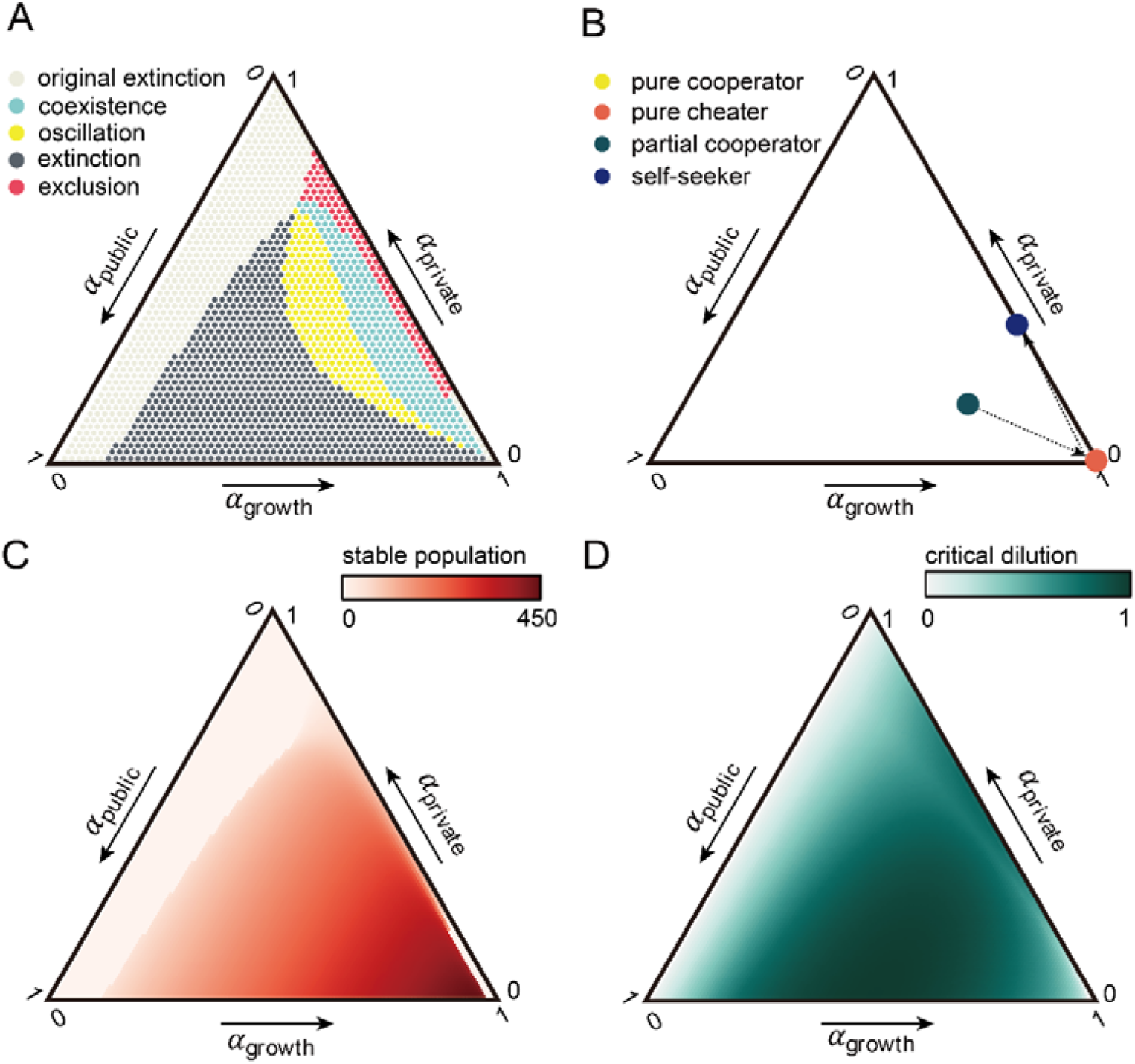
Assessment of all possible strategies, by their interplay with cheaters, evolutionary stability, stable population, and critical dilution rate. (A) How different strategies interact with the pure cheater. Each strategy's competition outcomes with the pure cheater are indicated by dot color in the ternary plot (light gray: non-viable by itself; deep gray: viable by itself but extinct with the cheater; yellow: oscillate with the cheater; blue: stably coexist with the cheater; red: exclude the cheater). (B) The chain of invasion in the strategy space. For each arrow, the dot at the beginning of the arrow is the initial strategy that creates the steady-state environment, and the dot at the endpoint of the arrow indicates the strategy with the maximal growth rate in the formal environment. The endpoint of the whole invasion chain is the evolutionarily stable strategy that can not be invaded by any other strategies. The path starts with a partial cooperator strategy(green dot, 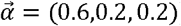), then directs to the pure cheater strategy(orange dot), and ends at the self-seeker strategy(blue dots, 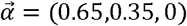). (C) The stable population for different strategies when existing alone. (D) The maximal dilution rate for non-zero biomass in a steady-state chemostat.

We evaluated a species’ “non-invasive strategy” by employing the invasion chain method(Taillefumier, Posfai et al. 2017): starting with one of the arbitrarily chosen strategies, we added strains with the highest growth rate in the existing chemical environment into the environment until no species could be added. A typical invasion chain is depicted in Figure 4B: the pure cheater has the highest growth rate in the environment shaped by the initial partial cooperator, but in the environment co-created by these two strains, the self-seekers (*α*_public_ = 0, *α*_private_ > 0) have a growth advantage, allowing them to invade and create an environment devoid of public siderophores. In such an environment, only self-seekers can survive, which brings the invasion chain to an end. In summary, within the same species, the self-seeker strategy is the most resistant against pure cheater strains and is “evolutionarily optimal”, because it generates a chemical environment that cannot be invaded by other strategies. Also, the self-seeker strategy can establish a community with a lower initial population, which benefits the start of colonization (Fig. S7).

However, when evaluating a species’ overall competitiveness, the self-seeker strategy is not always superior. For example, in evaluating the steady-state size of the population composed of a single strain (Fig. 4C), strategies that are close to the pure cheater have a larger overall population size. Meanwhile, in terms of resilience to harsher external environments, pure cooperators are more resistant to increased dilution (Fig. 4D).

Regarding the competition between species using different public siderophores, strains that invest more in public siderophores are more capable of invading another species (Fig. S4A), whereas strains that invest less in private siderophores are more resistant to invasion by another species (Fig. S4C). Self-seeker strategies, on the other hand, are not effective both in invasion and resistance(Fig. S4B, D).

## Discussion

Understanding diversity has long been a cornerstone of microbial ecology (Delmont, Robe et al. 2011). While the number of species may be limited by chemical dimensions, microorganisms are capable of expanding this limit by actively generating chemical diversity in their microhabitat. In this work, we modeled the specific system of siderophore-mediated iron competition in order to investigate more general questions: first, how could cooperators responsible for chemical diversity resist extinction induced by cheaters; and second, what is the ecological consequence of microbes creating new chemical dimensions. For the first question, we suggested the privatization of siderophores acting as a “game changer” to allow partial cooperators advantage over pure cheaters, and derived that a necessary condition for coexistence is that the cooperators who invest more in public siderophores also need to invest more in private siderophores. Concerning the second question, we analytically compared the difference in stability criteria between the traditional consumer resource models and the “resource partition model,” in which organisms not only consume but also produce resources. In addition, the public siderophore does not become a “resource” until it forms an association with ferric iron, the actual resource that microorganisms require. This is an additional layer of meaning for the term “resource-partition models”: externally supplied resources (iron) and microbe-generated resources (siderophore) interact to form the actual resources (iron-associated siderophores) taken up by the microorganisms.

Microbes interact via influencing their shared environment (McGill, Enquist et al. 2006). Classical ecological models emphasize the “consumption” aspect of such influences (Tikhonov and Monasson 2017, Altieri and Franz 2019). With the rapid development of microbiology, it has become increasingly apparent that bacteria have enormous potential for introducing new chemicals into their microhabitats(Gavriilidou, Kautsar et al. 2022). Siderophore is just one of the many secondary metabolites that microbes actively produce for their own benefit. Antibiotics, bacteriocins, signaling molecules, and even bacterial vesicles can all be considered “chemical dimensions” generated by microbes themselves (Bajić, Rebolleda-Gómez et al. 2021, Niehus, Oliveira et al. 2021). Regarding these “self-generated dimensions,” a general ecological framework has yet to be established. The conventional consumer resource model provides intuitive coexistence criteria (i.e. species should preferentially consume the resource that limits their growth), with consumption vectors clearly segregating supply conditions into zones of the same stability (Tilman 1982, Koffel, Daufresne et al. 2016). In this “resource partition model”, however, due to the reversed direction of consumption vectors, there are additional parameter areas where the RH criterion can change signs. Consequently, in the regions of steady-state equilibrium in the classical model, non-equilibrium dynamics become possible. In general, our analysis suggested that an ecosystem with “self-generated dimensions” tends to be more dynamic and complex.

Oscillation usually bridges two distinctive states and often plays critical roles in various biological systems (Goldbeter 2008). Notably, the oscillation in our model differs from the oscillation in the work of Huisman et al. where the oscillation bridges between the stable equilibrium and the chaos with higher-than-CEP dynamical coexistence (Huisman and Weissing 1999, Huisman and Weissing 2001). In our model, the parameter region of oscillation locates between total extinction and stable coexistence between partial cooperators and cheaters. Oscillation here is more of a “danger zone” indicating that the system is on the verge of collapse, similar to the early warning signatures of ecosystems (Carpenter, Cole et al. 2011). Indeed, our continuous equations assume that organisms can recover from arbitrarily small values, but the troughs of oscillatory dynamics increase the probability of stochastic extinction in real systems (Reichenbach, Mobilia et al. 2006, Ovaskainen and Meerson 2010). Recent studies in the field of game theory, however, have detected oscillations when the environment is made explicit and have shown that oscillatory dynamics prevent the extinction of cooperators (Weitz, Eksin et al. 2016, Hilbe, Šimsa et al. 2018, Tilman, Plotkin et al. 2020). Given the specificity of oscillatory dynamics in ecology and the propensity of resource partition models to enter the oscillation zone, it would be intriguing to explore the facilitative or destructive functions of oscillations in diverse ecological systems with chemical innovations.

In the specific game of microbial iron competition, the pervasiveness of siderophore production remains the subject of continuing investigation (Barber and Elde 2015, Kramer, Özkaya et al. 2020, Lee, Eldakar et al. 2021). Our work emphasized that private siderophores only provide possible solutions in some species, such as marine microbes or mycobacterium, where the siderophores can be modified into membrane-attached form.

Other kinds of siderophore privatization, such as keeping siderophores intracellularly for relieving oxidative stress, have been proposed to confer cheater resistance (Jin, Li et al. 2018). Actually, we hypothesize that the division of the iron resource by siderophores provides a universal mechanism for “resource privatization”: numerous siderophores exist in the natural world, each with their own specific receptors (Cornelis and Matthijs 2002, Jin, Li et al. 2018). For a given type of siderophore, microorganisms with corresponding receptors can share it as a public good, whereas microbes without comparable receptors perceive it as inaccessible “private goods” (Leventhal, Ackermann et al. 2019). Theoretical models suggested that the populations of different cooperators utilizing distinct siderophores can be regulated by their cheaters, similar to how parasites impose negative frequency selection (Lee, van Baalen et al. 2012). In addition, a model with different types of siderophores suggested that coexistence between cooperators and cheaters is possible if a “loner” uses a kind of relatively inefficient siderophores (Inglis, Biernaskie et al. 2016). In the future, it would be exciting to investigate iron interactions in a more realistic biological setting and systematically investigate the ecological consequences of microbial chemical innovation.

## Data Availability

The source code and parameters used are available in the supplementary material.

## Acknowledgements

This work was supported by grants from Peking-Tsinghua Center for Life Sciences.

## Author contributions

Jiqi Shao performed the majority of computational and mathematical analysis in this research and drafted the manuscript. Nan Rong performed the preliminary computational analysis in this research and drafted the manuscript. Zhenchao Wu, Shaohua Gu, Beibei Liu, and Ning Shen offered insightful comments and assisted in revising the manuscript. Zhiyuan Li conceptualized and oversaw the project and revised the manuscript. All authors gave final approval for publication and agreed to be held accountable for the work performed herein.

