## Supplementary Information for "Siderophore-mediated iron partition promotes dynamical coexistence between cooperators and cheaters"

### Supplemental Table 1: Symbols

| Chemostat parameters | |
| --- | --- |
| $R_{iron, supply}$ | Iron supply. |
| $d$ | Dilution rate. |
| Chemostat variables | |
| $R_{\mathrm{sid}}$ | Concentration of public siderophores. |
| $R_{\mathrm{iron}}$ | Concentration of the iron. |
| $m_{\sigma}$ | Biomass density of microbe $\sigma$. |
| species-specific quantities | |
| $\vec{\alpha}_{\sigma}=\left( \alpha_{\sigma, \mathrm{growth}},\alpha_{\sigma, \mathrm{private}},\alpha_{\sigma, \mathrm{public}} \right)$ | Resource allocation strategy of microbe $\sigma$, defined by the fractions of resources being located to growth ($\alpha_{\sigma, growth}$), production of private siderophores$\left( \alpha_{\sigma, private} \right)$, and production of public siderophores ($\alpha_{\sigma, public}$) |
| $g\left( R_{\mathrm{sid}},R_{\mathrm{iron}},\vec{\alpha}_{\sigma} \right)$ | Growth rate as a function of $R_{\mathrm{sid}},R_{\mathrm{iron}},$and $\vec{\alpha}_{\sigma}$. |
| $I_{\sigma}=I\left( R_{\mathrm{sid}},R_{\mathrm{iron}},\vec{\alpha}_{\sigma_{i}} \right)$ | Iron intake rate per biomass of the microbe $\sigma_{i}$. |
| $v_{m},v_{l}$ | Uptake coefficients for private and public siderophores, respectively. “m” stands for “membrane”, and “l” stands for “liquid” |
| $K_{m},K_{l}$ | Affinity constants for private and public siderophores, respectively |
| $\gamma$ | Species growth coefficient. |
| $\beta$ | Production coefficient of private siderophores. |
| $\epsilon$ | Production coefficient of public siderophores. |
| $p$ | Consumption ratio of public siderophores when microbes uptake iron. |
| $r$ | Relative biomass concentration in the cell compared to its microenvironment, set to be a constant of 100. |

### 1. Description of the resource partition model

We utilized the chemostat-typed model to investigate the ecological interactions between iron-uptake strategies. In this simplified iron-partition model, we made the following assumptions:

1. Iron is the primary growth-limiting resource, and the iron intaking rate linearly scales with the growth rate.
2. Siderophores can be recycled during the iron-uptake process, so in the equations, we excluded their consumption for simplicity. Actually, we also demonstrated that including siderophores consumption in the model does not qualitatively alter the results of this investigation (Figure S5).
3. “Species” in this work were represented by different parameters, such as $v_{m}$, $K_{l}$, $\gamma$, $\beta$, etc.; Within a species, different strains adopt different resource allocation strategies $\vec{\alpha}_{\sigma}=\left( \alpha_{\sigma, growth},\alpha_{\sigma, private},\alpha_{\sigma, public} \right)$, which specifies the fraction of cellular proteins and energies devoted to the growth, private siderophores production, and public siderophores production, respectively. To represent the limited cellular resources, the summation of all components in $\vec{\alpha}_{\sigma}$ equals to one. By this measurement, behaviors can be quantified by elements of $\vec{\alpha}_{\sigma}$. For example, “cheating” is defined as not producing public siderophores ($\vec{\alpha}_{\sigma, \mathrm{public}}=0$).

Thereby, a continuous spectrum of strategies could be visualized in a ternary diagram,

where we can define four major classes of strategies:

- Pure cheater: $\vec{\alpha}_{\sigma}=\left( 1,0,0 \right)$, where the strain does not produce any siderophore but allocates all resource budgets into growth.
- Self-seeker:$\vec{\alpha}_{\sigma}=\left( \alpha_{\sigma, growth},1-\alpha_{\sigma, growth},0 \right)$, where the strain only produces private siderophores without secreting them. Self-seekers can be considered as another type of cheater as it does not produce public siderophore but can utilize them.
- Pure cooperator: $\vec{\alpha}_{\sigma}=\left( \alpha_{\sigma, \mathrm{growth}},0,1-\alpha_{\sigma, growth} \right)$, where the strain only produces public siderophores without privatizing them.
- Partial cooperator: $\vec{\alpha}_{\sigma}=\left( \alpha_{\sigma, \mathrm{growth}},\alpha_{\sigma, private},\alpha_{\sigma, public} \right)$, with all three fractions non-zero. In this case, the strain produces both public and private siderophores.

Following the assumptions above, resource competition dynamics between two strains 1 and 2 adopting strategies $\vec{\alpha}_{1}$ and $\vec{\alpha}_{2}$ can be expressed as the following:

$$\frac{dm_{1}}{dt}=m_{1}\left( g\left( R_{\mathrm{iron}},R_{\mathrm{sid}},\vec{\alpha}_{1} \right)-d \right)$$

(1)

$$\frac{dm_{2}}{dt}=m_{2}\left( g\left( R_{\mathrm{iron}},R_{\mathrm{sid}},\vec{\alpha}_{2} \right)-d \right)$$

(2)

$$\frac{dR_{\mathrm{sid}}}{dt}=\frac{m_{1}}{r}\alpha_{\text{1, }\text{public}}\epsilon+\frac{m_{2}}{r}\alpha_{\text{2, }\text{public}}\epsilon-dR_{\mathrm{sid}}$$

(3)

$$\frac{dR_{\mathrm{iron}}}{dt}=d\left( R_{iron, supply}-R_{\mathrm{iron}} \right)-\sum_{i=1}^{2} \frac{m_{i}}{r}I\left( R_{\mathrm{iron}},R_{\mathrm{sid}},\vec{\alpha}_{i} \right)$$

(4)

The total iron intake rate is the summation of iron intake from both private and public sources:

$$I\left( R_{\mathrm{iron}},R_{\mathrm{sid}},\vec{\alpha}_{i} \right)=J_{\sigma_{i},\mathrm{private}}+J_{\sigma_{i}, public},$$

(5)

where $J_{\sigma,\mathrm{private}}$ and $J_{\sigma, \mathrm{public}}$ are the uptake fluxes of iron from private and from public siderophores, respectively, which we assumed Monad forms as:

$$J_{\sigma,\mathrm{private}}=\alpha_{\sigma,\mathrm{private}}\beta v_{m}\frac{R_{\mathrm{iron}}}{K_{m}+R_{\mathrm{iron}}},$$

(6)

$$J_{\sigma, \mathrm{public}}=v_{l}R_{\mathrm{sid}}\frac{R_{\mathrm{iron}}}{K_{l}+R_{\mathrm{iron}}}.$$

(7)

$v_{m}$ and $v_{l}$ denote the intake rate coefficients, and we set $v_{m}=v_{l}=1$ for convenience. $\beta$ represents the efficiency of private siderophores’ production. $K_{m}$ and $K_{l}$ are the affinity constants for private siderophores and public siderophores. In this model, we set $K_{m}>K_{l}$ to show that public siderophores have advantages in acquiring iron. In the later section, we will prove that other forms of iron uptake yield to similar conclusions.

By the first assumption, the growth rate linearly scales with the total iron intake $I$, as well as the fraction of resources allocated to biomass accumulation $\alpha_{\sigma, \mathrm{growth}}$:

$$g\left( R_{\mathrm{iron}},R_{sid},\vec{\alpha}_{\sigma} \right)=\gamma\alpha_{\sigma, \mathrm{growth}}I\left( R_{\mathrm{iron}},R_{\mathrm{sid}},\vec{\alpha}_{\sigma} \right).$$

(8)

### 2. Conditions for the stable coexistence between two strains

#### 2.1.  Growth contour for a single strain in the chemical space

In this work, the two-dimensional “chemical space” is defined by the concentrations of iron and siderophores that directly impact cellular growth, as the ensembles of all possible $\left[ R_{\mathrm{iron}},R_{\mathrm{sid}} \right]$.

The growth contour of strain $\sigma$ is defined as all $\left[ R_{\mathrm{iron}},R_{\mathrm{sid}} \right]$ that meet $g\left( R_{\mathrm{iron}},R_{sid},\vec{\alpha}_{\sigma} \right)=d$ in the chemical space, equivalent to the zero net growth isoclines (ZNGI) in contemporary niche theory. For strain $\sigma$, we can obtain the growth contour from Eq.(1) equals 0, leading to the following relation between $R_{\mathrm{iron}}$ and $R_{\mathrm{sid}}$ for the growth contour:

$$R_{\mathrm{sid}}=\frac{\frac{d}{\gamma\alpha_{\sigma, \mathrm{growth}}}-\frac{v_{m}{\beta\alpha}_{\sigma,\mathrm{private}}R_{\mathrm{iron}}}{K_{m}+R_{\mathrm{iron}}}}{\frac{v_{l}R_{\mathrm{iron}}}{K_{l}+R_{\mathrm{iron}}}}$$

(9)

#### 2.2.  Requirement for the presence of intersection between two growth contours

For multiple strains, the possible chemical environment allowing for stable coexistence must locate at the intersection of their growth contours, indicated by the “*”.

Between two strains 1 and 2 adopting two strategies, the relationships between the intersection point $R_{\mathrm{iron}}^{*}$ and $R_{\mathrm{sid}}^{*}$ follows:

$$R_{\mathrm{sid}}^{*}=\frac{v_{m}\beta\left( K_{l}+R_{\mathrm{iron}}^{*} \right)\left( \alpha_{1, \mathrm{growth}}\alpha_{1,\mathrm{private}}-\alpha_{2, \mathrm{growth}}\alpha_{2, \mathrm{private}} \right)}{v_{l}\left( K_{m}+R_{\mathrm{iron}}^{*} \right)\left( \alpha_{2, \mathrm{growth}}-\alpha_{1, \mathrm{growth}} \right)}.$$

(10)

Given a certain dilution rate $d$, $R_{\mathrm{iron}}^{*}$ and $R_{\mathrm{sid}}^{*}$ could be solved as a function of $d$:

$$R_{\mathrm{iron}}^{*}=\frac{-dK_{m} \left( \alpha_{1, \mathrm{growth}}-\alpha_{2, \mathrm{growth}} \right)}{d\left( \alpha_{1, \mathrm{growth}}-\alpha_{2, \mathrm{growth}} \right)+\beta\gamma v_{m}\alpha_{1, \mathrm{growth}}\alpha_{2, \mathrm{growth}}\left( \alpha_{1,\mathrm{private}}-\alpha_{2, \mathrm{private}} \right)}$$

(11)

$$R_{\mathrm{sid}}^{*}=-\left( \alpha_{1, \text{growth}}\alpha_{1,\text{private}}-\alpha_{2, \text{growth}}\alpha_{2, \text{private}} \right)\left( K_{l}\left( \alpha_{1, \text{growth}}\left( \beta\gamma v_{m}\alpha_{2, \text{growth}}\left( \alpha_{1,\text{private}}-\alpha_{2, \text{private}} \right)+d \right)-d\alpha_{2, \text{growth}} \right)+{dK}_{m}\left( \alpha_{2, \text{growth}}-\alpha_{1, \text{growth}} \right) \right)\left( \gamma v_{l}K_{m}\alpha_{1, \text{growth}}\alpha_{2, \text{growth}}\left( \alpha_{1, \text{growth}}-\alpha_{2, \text{growth}} \right)\left( \alpha_{1, \text{private}}-\alpha_{2, \text{private}} \right) \right)^{-1}$$

(12)

From the solution and trajectory derived above, the presence of intersection points requires:

$\alpha_{1, \text{growth}}\neq\alpha_{2, \text{growth}} , \alpha_{1,\text{private}}\neq\alpha_{2, \text{private}}$*,*

(13)

so that the denominators of Eq. (11)-(12) can be non-zero.

In the following analysis, we always assume that strategy 1 allocate less on its own growth than strategy 2, then we have the setting:

$$\alpha_{2, \mathrm{growth}}-\alpha_{1, \mathrm{growth}}>0.$$

(14)

If the intersection point for possible coexistence exists with realistic value ($R_{\mathrm{iron}}^{*}>0,R_{\mathrm{sid}}^{*} >0$), then the numerator of Eq.(10) needs to be larger than zero too:

$$\alpha_{1, \mathrm{growth}}\alpha_{1,\mathrm{private}}-\alpha_{2, \mathrm{growth}}\alpha_{2, \mathrm{private}}>0.$$

(15)

Eq.(15) is the condition for the presence of the intersection point.

By Eq.(15) and with the setting $\alpha_{2,\mathrm{growth}}-\alpha_{1,\mathrm{growth}}>0$., it is straightforward to conclude the extinction of the pure cooperator (strain 1) when conferring cheaters (strain 2): for a pure cooperator, $\alpha_{1,\mathrm{private}}=0$, therefore $\alpha_{1,\mathrm{growth}}\alpha_{1,\mathrm{private}}-\alpha_{2,\mathrm{growth}}\alpha_{2,\mathrm{private}}<0$ for any none zero $\alpha_{2,\mathrm{private}}$. Therefore, coexistence is not possible for a pure cooperator with any other types of strategies, let alone cheaters.

#### 2.3.  Requirement for the presence of intersection between a partial cooperator and a pure cheater

If we set strategy 1 being a partial cooperator ($\alpha_{1,\mathrm{private}}>0$, $\alpha_{1, \mathrm{public}}>0$) and strategy 2 being a pure cheater$(\alpha_{2, \mathrm{growth}}=1$), according to Eq.(11)-(15), the necessary condition for a positive $R_{\mathrm{iron}}^{*}$ is:

$$\alpha_{1,\mathrm{private}}-\alpha_{2, \mathrm{private}}>0$$

(16)

Therefore, $\alpha_{1, \mathrm{growth}}<\alpha_{2, \mathrm{growth}}$, $\alpha_{1,\mathrm{private}}>\alpha_{2, \mathrm{private}}$ is required for the intersection point to exist, therefore is the necessary condition for coexistence. This requirement means that only the partial cooperators ($\alpha_{1,\mathrm{private}}>0$) can possibly stably coexist with the pure cheater ($\alpha_{2, \mathrm{growth}}=1$), while the pure cooperator goes to extinction when the cheater invades. More generally, for the possible coexistence between any pair of strains, the one that allocates fewer resources in growth must invest more in private siderophores.

#### 2.4. Requirement for non-zero biomass of both strains

Stable coexistence between two strains also requires the positive biomass of both strains. At the possible steady state of coexistence, we can set Eq.(3)-(4) equal to 0. Then, the solutions for $m_{1}^{*}$ and $m_{1}^{*}$ can be deduced as a function of the steady state concentrations of the iron $R_{\mathrm{iron}}^{*}$ and the siderophore $R_{\mathrm{sid}}^{*}$:

$$m_{1}^{*}=r\frac{\frac{d}{\epsilon}\frac{R_{\mathrm{sid}}^{*}}{\alpha_{2, \mathrm{growth}}}-\gamma\alpha_{2\text{, }\text{public}}\left( R_{iron, supply}-R_{\mathrm{iron}}^{*} \right)}{\frac{\alpha_{\text{1, }\text{public}}}{\alpha_{2, \mathrm{growth}}}-\frac{\alpha_{\text{2, }\text{public}}}{\alpha_{1, \mathrm{growth}}}},$$

(17)

$$m_{2}^{*}=r\frac{\gamma\alpha_{\text{1, }\text{public}}\left( R_{iron, supply}-R_{\mathrm{iron}}^{*} \right)-\frac{d}{\epsilon}\frac{R_{\mathrm{sid}}^{*}}{\alpha_{1, \mathrm{growth}}}}{\frac{\alpha_{\text{1, }\text{public}}}{\alpha_{2, \mathrm{growth}}}-\frac{\alpha_{\text{2, }\text{public}}}{\alpha_{1, \mathrm{growth}}}}.$$

(18)

For stable coexistence, $m_{1}^{*}$ and $m_{2}^{*}$ need to be positive.

As will be proved by the next section, under the setting that $\alpha_{1, \mathrm{growth}}<\alpha_{2, \mathrm{growth}}$.the denominator of the formal two equations, $\frac{\alpha_{\text{1, }\text{public}}}{\alpha_{2, \mathrm{growth}}}-\frac{\alpha_{\text{2, }\text{public}}}{\alpha_{1, \mathrm{growth}}}>0$ for stable coexistence.

Therefore, in order for $m_{1}^{*}>0$, it requires that $\frac{d}{\epsilon}\frac{R_{\mathrm{sid}}^{*}}{\alpha_{2, \mathrm{growth}}}-\gamma\alpha_{2\text{, }\text{public}}\left( R_{iron, supply}-R_{\mathrm{iron}}^{*} \right)>0$. Similarly, $m_{2}^{*}>0$ requires$\gamma\alpha_{\text{1, }\text{public}}\left( R_{iron, supply}-R_{\mathrm{iron}}^{*} \right)-\frac{d}{\epsilon}\frac{R_{\mathrm{sid}}^{*}}{\alpha_{1, \mathrm{growth}}}>0$. Together, they lead to the requirement of supply concentration that:

$R_{\mathrm{iron}}^{*}+\frac{1}{k_{2}}R_{\mathrm{sid}}^{*}>R_{iron, supply}>R_{\mathrm{iron}}^{*}+\frac{1}{k_{1}}R_{\mathrm{sid}}^{*}$,

(19)

where $k_{1}=\frac{\epsilon\gamma}{d}\alpha_{1, \mathrm{public}}\alpha_{1, \mathrm{growth}}$ , $k_{2}=\frac{\epsilon\gamma}{d}\alpha_{2, \mathrm{public}}\alpha_{2, \mathrm{growth}}$.

In these graphical tools developed by Tilman et al. ^1^, The consumption vector is defined as $\vec{C_{i}}=\left[ \begin{aligned} \partial_{m_{i}}\overset{.}{R_{1}} \\ \partial_{m_{i}}\overset{.}{R_{2}} \end{aligned} \right]$ for species $i$, whose slope indicates the consumption preference of species $i$. $k_{1}$ and $k_{2}$ are the slopes of the consumption vectors of strain 1 and strain 2, respectively. and $R_{\mathrm{iron}}^{*}+\frac{1}{k_{i}}R_{\mathrm{sid}}^{*}$ is equal to the intersection of the reverse extension of the consumption vector of strain $i$ with the x-axis.

Taken together, $m_{1}^{*},m_{2}^{*}>0$ require the iron supply point, $R_{iron, supply},$locates inside the sector spanned by the inverse extensions of supply vectors of two strains, just as the classical consumer resource model.

Nevertheless, the presence of growth contour intersection, and non-zero biomass of two strains at the intersection point, only provide necessary conditions for the coexistence. The stability of the possible coexistence points needs to be analyzed by the Jacobian matrix.

### 3. Analysis of the stability of coexistence and the consequences of the self-creating dimension

Unlike the classical consumer resource model where species utilizes externally-supplied resources, microbe in our model can access external iron resources more efficiently by producing public siderophores. In terms of chemical dimensions, the production of public goods implies that bacteria autonomously generate new resource dimensions (such as $\left[ R_{sid} \right]$ in our model). This resource-generation behavior changes the sign of the consumption vector, and can further affect the stability criteria for the system compared to the classical consumer resource models.

In the following section, we first reviewed the classical consumer resource model (Tilman’s model) and its stability criteria, and further elucidated the updated stability criteria in the resource partition model, using the iron-partition model in Eq. (1)-(4) as example.

#### 3.1.  The general form of resource partition model and its stability criteria

Similar to a classical consumer resource model (represented by Tilman’s model), the generalized form of microbes interacting with two resources $R_{1}$ and $R_{2}$ can be represented as:

$$\frac{dm_{i}}{dt}=m_{i}\left( g_{i}\left( R_{1},R_{2} \right)-d \right),\text{ for }i=1,2$$

(20)

$$\frac{dR_{j}}{dt}=d\left( R_{j, \mathrm{supply}}-R_{j} \right)-\sum_{i=1}^{2} m_{i}h_{ij}\left( R_{1},R_{2} \right)g_{i}\left( R_{1},R_{2} \right),\text{ for }j=1,2,$$

(21)

Where $R_{j, \mathrm{supply}}$ is the supply concentration of resource $j$, and $h_{ij}$ is the function describing the amount of resource $j$ impacted by species $i$ per-biomass.

When $h_{ij}$ describes resource uptake and has a positive sign, Eq. (20)-(21) above represent the classical consumer resource model^1^. When at least one of the $h_{ij}$ describes resource production and exhibit negative sign, these equations represents the broader “resource partition model” and exhibit different criteria on the stability of coexistence.

The graphical approach is composed of two elements: the growth contour, i.e. the zero net growth isoclines (ZNGI), and the consumption vector. ZNGI, i.e. the growth contour, is defined as all $\left[ R_{1},R_{2} \right]$ that meet $g\left( R_{1},R_{2} \right)=d$. And consumption vectors  $\vec{C_{i}}=-d\left[ \begin{aligned} h_{i1} \\ h_{i2} \end{aligned} \right]$ indicates the total consumption rate of resources per biomass of the species $i$ at the growth contour.

Assume that the growth contours of two species intersect at a fixed point $\left[ R_{1}^{*},R_{2}^{*} \right]$. The resource supply vector $\vec{U}=d\left[ \begin{aligned} R_{1, \mathrm{supply}}-R_{1}^{*} \\ R_{2, supply}-R_{2}^{*} \end{aligned} \right]$ points to the supply point $\left[ R_{1, supply}, R_{2, supply} \right]$ from the fixed point. To balance $\vec{U}$, the summation of resource supply vector $\vec{U}$ and consumption vectors  $\vec{C_{1}}$ and  $\vec{C_{2}}$ equals to $0$:$m_{1}^{*}\vec{C_{1}}+m_{2}^{*}\vec{C_{2}}+\vec{U}=\vec{0}$.

For the fixed point to exist with non-zero biomass of strain 1 and 2, the resource supply vector needs to be located within the sector area formed by two consumption vectors $\vec{C_{1}}$ and $\vec{C_{2}}$. Nevertheless, the (1) existence of the intersection of two growth contours, and (2) supply vector locates within the angle of consumption vectors, are only necessary conditions for coexistence. The stability of coexistence needs to also be accessed by the eigenvalue of the Jacobian matrix of the fixed point.

At the fixed point (*), the Jacobian matrix of the models above takes a general form:

$$J= \left[ \begin{matrix} 0 & 0 & v_{11} & v_{12} \\ 0 & 0 & v_{21} & v_{22} \\ -w_{11} & -w_{12} & -x_{11} & -x_{12} \\ -w_{21} & -w_{22} & -x_{21} & -x_{22} \end{matrix} \right] ,$$

(22)

with

$$v{}_{ij}=\left( \frac{\partial\overset{.}{m_{i}}}{\overset{.}{\partial R_{j}}} \right)^{*} , w{}_{ij}=-\left( \frac{\partial\overset{.}{R_{i}}}{\overset{.}{\partial m_{j}}} \right)^{*},x{}_{ij}=-\left( \frac{\partial\overset{.}{R_{i}}}{\overset{.}{\partial R_{j}}} \right)^{*}.$$

For simplicity, we set the abbreviation as:

$$P_{1}=\left( \partial_{R_{1}}g_{1} \right)^{*} , P_{2}=\left( \partial_{R_{2}}g_{1} \right)^{*},$$

$$P_{3}=\left( \partial_{R_{1}}g_{2} \right)^{*} , P_{4}=\left( \partial_{R_{2}}g_{2} \right)^{*},$$

$$c_{11}=\left( h_{11} \right)^{*}, c_{12}=\left( h_{21} \right)^{*},$$

$$c_{21}=\left( h_{12} \right)^{*}, c_{22}=\left( h_{22} \right)^{*},$$

(23)

so we have elements in the Jacobian matrix expressed as:

$$v_{11}=m_{1}^{*}P_{1}, v_{12}=m_{1}^{*}P_{2},$$

$$v_{21}=m_{2}^{*}P_{3}, v_{22}=m_{2}^{*}P_{4}$$

$$w_{11}=c_{11}d, w_{12}=c_{12}d,$$

$$w_{21}=c_{21}d, w_{22}=c_{22}d.$$

$$x_{11}=d,\text{ }x_{12}=0,$$

$$x_{21}=m_{1}^{*}c_{21}P_{1}+m_{2}^{*}c_{22}P_{3},\text{ }x_{22}=d+m_{1}^{*}c_{21}P_{2}+m_{2}^{*}c_{22}P_{4}$$

(24)

At the steady state, ${g_{1}}^{*}={g_{2}}^{*}=d$. With definitions in Eq.(23)-(24), the characteristic equation of eigenvalue $\lambda$ for the Jacobian matrix can be expressed as:

$$\lambda^{4}+\left( x_{11}+x_{22} \right)\lambda^{3}+\left( q_{1}+q_{4}+x_{11}x_{22}-x_{12}x_{21} \right)\lambda^{2}+\left( x_{11}q_{4}+x_{22}q_{1}-x_{12}q_{3}-x_{21}q_{2} \right)\lambda+\left( q_{1}q_{4}-q_{2}q_{3} \right)=0,$$

(25)

the coefficients of this quartic equation are: $a_{0}=1$, $a_{1}=x_{11}+x_{22}$, $a_{2}=q_{1}+q_{4}-x_{12}x_{21}+x_{11}x_{22},$ $a_{3}=q_{4}x_{11}-q_{3}x_{12}-q_{2}x_{21}+q_{1}x_{22}$, $a_{4}=-q_{2}q_{3}+q_{1}q_{4}$, with $q_{i}$ defined as:

$$q_{1}=w_{11}v_{11}+w_{12}v_{21},$$

$$q_{2}=w_{11}v_{12}+w_{12}v_{22},$$

$$q_{3}=w_{21}v_{11}+w_{22}v_{21},$$

$$q_{4}=w_{21}v_{12}+w_{22}v_{22}.$$

(26)

The Routh-Hurwitz (RH) criterion states a necessary and sufficient condition for the stability of a dynamical system ^2^: for a quartic equation for the eigenvalue $\lambda$, $a_{0}\lambda^{4}+a_{1}\lambda^{3}+a_{2}\lambda^{2}+a_{3}\lambda+a_{4}=0$, The RH table were calculated as Supplemental Table 2:

### Supplemental Table 2: The Routh-Hurwitz table

| Term ID | Values |
| --- | --- |
| Term 0 | $a_{0}$ |
| Term 1 | $a_{1}$ |
| Term 2 | $\frac{a_{1}a_{2}-a_{0}a_{3}}{a_{1}}$ |
| Term 3 | $\frac{\left( a_{1}a_{2}-a_{0}a_{3} \right)a_{3}-a_{1}^{2}a_{4}}{a_{1}a_{2}-a_{0}a_{3}}$ |
| Term 4 | $a_{4}$ |

Along the “Values” column of Table S2, from term 0 to term 4, the number of sign changes equals the number of non-negative roots in Eq. (24). As $a_{0}=1$, the RH table leads to four RH criteria: (1) $a_{1}>0$, (2) $a_{4}>0$, (3) $a_{3}>0$, (4) $a_{1}a_{2}a_{3}-a_{1}^{2}a_{4}-a_{3}^{2}>0$. Detailed derivations of the RH criterion on resource competition models can be found in the appendix of Tilman’s original literature ^1^.

In the following text, we will go through these four RH criteria, and compare the difference between the classical consumer resource model and the resource partition model. We use the iron-partition model in Eq. (1)-(4) as examples, which specifies the parameters in Eq.(23) into:

$$P_{1}=\left( \partial_{R_{\mathrm{sid}}}g_{1} \right)^{*} , P_{2}=\left( \partial_{R_{\mathrm{iron}}}g_{1} \right)^{*},$$

$$P_{3}=\left( \partial_{R_{\mathrm{sid}}}g_{2} \right)^{*} , P_{4}=\left( \partial_{R_{\mathrm{iron}}}g_{2} \right)^{*}.$$

$$c_{11}=-\alpha_{1, \mathrm{public}}\epsilon/r, c_{12}=-\alpha_{2, \mathrm{public}}\epsilon/r,$$

$$c_{21}=1/\left( \alpha_{1, \mathrm{growth}}\gamma r \right), c_{22}=1/\left( \alpha_{2, \mathrm{growth}}\gamma r \right).$$

(27)

Note that $c_{11},c_{12}<0$ here, which is different from the classical consumer resource model, as the public siderophore were produced instead of consumed by microbes.

#### 3.2.  Comparison of the four RH criteria between the classical and the iron-partition model

##### Criteria 1: $\boldsymbol{a}_{\boldsymbol{1}}\boldsymbol{>0}$

With the corresponding parameters replaced by definitions in Eq.(23)-(25), $a_{1}=x_{11}+x_{22}$ turns into:

$$a_{1}=2d+c_{21}P_{2}m_{1}^{*}+c_{22}P_{4}m_{2}^{*}.$$

(28)

In both classical model and the iron-partition model, $P$ denotes the growth rate dependency on “resources” and is always positive. Because all the terms on the right side are greater than or equal to zero, $a_{1}>0$ holds for both classical consumer resource model and the iron-partition model (Eq. (1)-(4)).

##### Criteria 2: $\boldsymbol{a}_{\mathbf{4}}\mathbf{>0}$

We discussed $a_{4}$ before $a_{3}$, as $a_{4}$ composed the second term of $a_{3}$. In the classical model, if criteria (2) $a_{4}>0$, then the criteria (3) $a_{3}>0$ holds automatically.

In both consumer resource and resource partition models, by definitions in Eq. (25), the original form $a_{4}=-q_{2}q_{3}+q_{1}q_{4}$ expand into:

$$a_{4}=dm_{1}^{*}m_{2}^{*}\left( c_{12}c_{21}-c_{11}c_{22} \right)\left( P_{2}P_{3}-P_{1}P_{4} \right),$$

(29)

There are two possible situations for $\left( c_{12}c_{21}-c_{11}c_{22} \right)\left( P_{2}P_{3}-P_{1}P_{4} \right)>0$:

| situation 1: | $\frac{P_{2}}{P_{1}}>\frac{P_{4}}{P_{3}} ,\frac{c_{11}}{c_{21}}<\frac{c_{12}}{c_{22}},$ | (30) |
| --- | --- | --- |
| or, situation 2 | $\frac{P_{2}}{P_{1}}<\frac{P_{4}}{P_{3}} ,\frac{c_{11}}{c_{21}}>\frac{c_{12}}{c_{22}}.$ | (31) |

The above requirements hold for both classical consumer resource and the iron-partition model. Nevertheless, these two types of models interpret the requirements differently.

In the classical consumer resource model, for situation 1, $\frac{P_{1}}{P_{2}}>\frac{P_{3}}{P_{4}}$ means the growth of species 1 is more limited by resource 1 compared to species 2, and $\frac{c_{11}}{c_{21}}>\frac{c_{12}}{c_{22}}$ means species 1 also preferentially consumes resource 1 compared to species 2. Similarly, for situation 2, the growth of species 1 is more limited by resource 2, and also consumes more of resource 2. In combination, both possibilities suggest that a necessary condition for stable coexistence in the classical consumer resource model is that each species must consume more of the one resource which more limits its own growth rate.

This condition can be easily applied by the graphical tool for judging coexistence (Figure S1). When the supply point locates within the region bounded by the reverse extensions of the consumption vectors, the growth contours in Figure S1 show the example that species A and species B are more limited by $R_{2}$ and $R_{1}$, respectively. As the consumption vectors show that species A and species B preferentially consume more $R_{1}$ and $R_{2}$ respectively, species A and B can stably coexist.

In the iron-partition model, we assumed $\alpha_{1, \mathrm{growth}}<\alpha_{2, \mathrm{growth}}$. If situation 1 holds, $\frac{P_{2}}{P_{1}}>\frac{P_{4}}{P_{3}}$ means the growth of strain 1, relative to strain 2, is more limited by iron. Since two strains have the same influx rate through public siderophores, $J_{12}=J_{22}$ and $\frac{\partial\left( J_{11}+J_{12} \right)}{\partial R_{\mathrm{sid}}}=\frac{\partial\left( J_{21}+J_{22} \right)}{\partial R_{\mathrm{sid}}}$. $\frac{P_{2}}{P_{1}}>\frac{P_{4}}{P_{3}}$, i.e., $\frac{\frac{\partial\left( J_{11}+J_{12} \right)}{\partial R_{\mathrm{iron}}}}{\frac{\partial\left( J_{11}+J_{12} \right)}{\partial R_{\mathrm{sid}}}}>\frac{\frac{\partial\left( J_{21}+J_{22} \right)}{\partial R_{\mathrm{iron}}}}{\frac{\partial\left( J_{21}+J_{22} \right)}{\partial R_{\mathrm{sid}}}}$, further demands $J_{11}>J_{21}$, i.e., ${\alpha_{1,private}>\alpha}_{2, \mathrm{private}}$. Note that $c_{11},c_{12}<0$, $\frac{c_{11}}{c_{21}}<\frac{c_{12}}{c_{22}}$ means $\left| \frac{c_{11}}{c_{21}} \right|>\left| \frac{c_{12}}{c_{22}} \right|$, that is, strain 1, relative to strain 2, produces proportionately more public siderophores. And $c_{12}c_{21}-c_{11}c_{22}=\frac{\epsilon\left( \frac{\alpha_{1, \mathrm{public}}}{\alpha_{2, \mathrm{growth}}}-\frac{\alpha_{2, \mathrm{public}}}{\alpha_{1, \mathrm{growth}}} \right)}{r^{2}\gamma}>0$ asks for $\alpha_{1, \mathrm{growth}}\alpha_{1, \mathrm{public}}-\alpha_{2, \mathrm{growth}}\alpha_{2, \mathrm{public}}>0$. Taken together, if the intersection point exists and situation 1 holds, under the assumption of

$${\alpha_{1, \text{growth}}<\alpha}_{2, \text{growth}},$$

(32)

$a_{4}>0$ requires that**:**

$${\alpha_{1,\text{private}}>\alpha}_{2, \text{private}},$$

(33)

$${\alpha_{1, \text{public}}>\alpha}_{2, \text{public}}\geq0.$$

(34)

If situation 2 holds and the assumption of $\alpha_{1, \mathrm{growth}}<\alpha_{2, \mathrm{growth}}$ applies, $\frac{P_{2}}{P_{1}}<\frac{P_{4}}{P_{3}}$ requires $J_{11}>J_{21}$, i.e., ${\alpha_{1,private}<\alpha}_{2, \mathrm{private}}$, which does not match the condition for the existence of intersection point ${\alpha_{1,private}>\alpha}_{2, \mathrm{private}}$. This would prove that situation 2 does not hold under the setting $\alpha_{1, \mathrm{growth}}<\alpha_{2, \mathrm{growth}}$.

Taken together, in the iron-partition model, if two strains could stably coexist, the strain that gives more public goods and plays a more cooperative role (strain 1) must retain more private goods than the other strain (strain 2).

##### Criteria 3: $\boldsymbol{a}_{\mathbf{3}}\mathbf{>0}$

With the corresponding items replaced by definitions in Eq.(23)-Eq.(25), the formula of $a_{3}$ turns into:

$$a_{3}=d\left( m_{1}^{*}\left( c_{11}P_{1}+c_{21}dP_{2} \right)+m_{2}^{*}\left( c_{12}P_{3}+c_{22}dP_{4} \right) \right)+m_{1}^{*}m_{2}^{*}\left( c_{12}c_{21}-c_{11}c_{22} \right)\left( P_{2}P_{3}-P_{1}P_{4} \right).$$

(35)

The second term is linear with $a_{4}$ in criteria (2).

In the classical consumer resource model, the resource is supplied externally and consumed by species, thus each of the $c_{11},c_{12},c_{21},c_{22}>0$. The first term in the equation above, $d\left( m_{1}^{*}\left( c_{11}P_{1}+c_{21}dP_{2} \right)+m_{2}^{*}\left( c_{12}P_{3}+c_{22}dP_{4} \right) \right)$, is non-negative under the classical model. Thus, as long as criteria (2) $a_{4}>0$ is satisfied, $a_{3}>0$ holds.

In the iron-partition model, however, the first term $d\left( m_{1}^{*}\left( c_{11}P_{1}+c_{21}dP_{2} \right)+m_{2}^{*}\left( c_{12}P_{3}+c_{22}dP_{4} \right) \right)$ is no longer guaranteed to be nonnegative, as $c_{11},c_{12}<0$. Therefore, the sign of $a_{3}$ could be negative even when the second term is positive (demonstrated by Region 2 in Figure 3B), adding a region of instability.

##### Criteria 4: $\boldsymbol{a}_{\mathbf{1}}\boldsymbol{a}_{\mathbf{2}}\boldsymbol{a}_{\mathbf{3}}\mathbf{-}\boldsymbol{a}_{\mathbf{1}}^{\mathbf{2}}\boldsymbol{a}_{\mathbf{4}}\mathbf{-}\boldsymbol{a}_{\mathbf{3}}^{\mathbf{2}}\mathbf{>0}$

After expansion and simplification, the equation in criteria (4) can be derived into the product of two terms:

$$a_{1}a_{2}a_{3}-a_{1}^{2}a_{4}-a_{3}^{2}=\left( 2d^{3}+dm_{1}^{*}\left( c_{11}P_{1}+2dc_{21}P_{2} \right)+dm_{2}^{*}\left( c_{12}P_{3}+2dc_{22}P_{4} \right)+m_{1}^{*}m_{2}^{*}\left( c_{12}c_{21}-c_{11}c_{22} \right)\left( P_{2}P_{3}-P_{1}P_{4} \right) \right)\left( \left( c_{11}P_{1}+dc_{21}P_{2} \right)m_{1}^{*}\left( d+c_{21}P_{2}m_{1}^{*} \right)+\left( c_{12}\left( dP_{3}+c_{21}P_{1}P_{4}m_{1}^{*} \right)+c_{22}\left( d^{2}P_{4}+P_{2}\left( c_{11}P_{3}+2dc_{21}P_{4} \right)m_{1}^{*} \right) \right)m_{2}^{*}+c_{22}P_{4}\left( c_{12}P_{3}+dc_{22}P_{4} \right){m_{1}^{*}}^{2} \right).$$

(36)

The first multiplier term $2d^{3}+dm_{1}^{*}\left( c_{11}P_{1}+2dc_{21}P_{2} \right)+dm_{2}^{*}\left( c_{12}P_{3}+2dc_{22}P_{4} \right)+m_{1}^{*}m_{2}^{*}\left( c_{12}c_{21}-c_{11}c_{22} \right)\left( P_{2}P_{3}-P_{1}P_{4} \right)$ is a linear combination of $a_{3}$ and $a_{4}$, which is non-negative when criteria (2) $a_{4}>0$ and criteria (3) $a_{3}>0$ holds.

Nevertheless, the second multiplier term differs in two types of models.

In the classical consumer resource model, with $c_{21},c_{22}>0$, the second multiplier term in Eq. (34), $\left( c_{11}P_{1}+dc_{21}P_{2} \right)m_{1}^{*}\left( d+c_{21}P_{2}m_{1}^{*} \right)+\left( c_{12}\left( dP_{3}+c_{21}P_{1}P_{4}m_{1}^{*} \right)+c_{22}\left( d^{2}P_{4}+P_{2}\left( c_{11}P_{3}+2dc_{21}P_{4} \right)m_{1}^{*} \right) \right)m_{2}^{*}+c_{22}P_{4}\left( c_{12}P_{3}+dc_{22}P_{4} \right){m_{1}^{*}}^{2}$ is always non-negative.

In the iron-partition model, with changed signs in $c_{11}$ and $c_{12}$, the sign of the second term $\left( c_{11}P_{1}+dc_{21}P_{2} \right)m_{1}^{*}\left( d+c_{21}P_{2}m_{1}^{*} \right)+\left( c_{12}\left( dP_{3}+c_{21}P_{1}P_{4}m_{1}^{*} \right)+c_{22}\left( d^{2}P_{4}+P_{2}\left( c_{11}P_{3}+2dc_{21}P_{4} \right)m_{1}^{*} \right) \right)m_{2}^{*}+c_{22}P_{4}\left( c_{12}P_{3}+dc_{22}P_{4} \right){m_{1}^{*}}^{2}$ cannot be indirectly inferred, and can still be negative even when the criteria (2) $a_{4}>0$ and criteria (3) $a_{3}>0$ holds, which contributes to the oscillatory dynamics (Region 3 in Figure 3, and Figure S3).

#### 3.3.  Summary of the conditions for stable coexistence in the iron-partition model.

First, in the iron-partition model of Eq.(1)-(4), the presence of the co-existence point between two strains, with the setting

$$\alpha_{1, \mathrm{growth}}<\alpha_{2, \mathrm{growth}},$$

requires

$$\alpha_{1,\mathrm{private}}>\alpha_{2, \mathrm{private}}.$$

Second, the non-zero biomass of two strains imposes constraints on the iron supply concentration $R_{iron, supply}$, that

$$R_{\mathrm{iron}}^{*}+\frac{1}{k_{2}}R_{\mathrm{sid}}^{*}>R_{iron, supply}>R_{\mathrm{iron}}^{*}+\frac{1}{k_{1}}R_{\mathrm{sid}}^{*}$$

with $k_{1}=\frac{\epsilon\gamma}{d}\alpha_{1, \mathrm{public}}\alpha_{1, \mathrm{growth}}$ , $k_{2}=\frac{\epsilon\gamma}{d}\alpha_{2, \mathrm{public}}\alpha_{2, \mathrm{growth}}$.

Graphically, this inequality requires the supply point [$R_{iron, supply}$, 0] locating within the region bounded by the reverse extensions of the consumption vectors in the chemical space.

Third, the stability of the coexistence point is judged by the RH criterion. The criteria (2) $a_{4}>0$ also requires

$${\alpha_{1, \text{public}}>\alpha}_{2, \text{public}}.$$

If it were a classical consumer resource model, the satisfaction of the criteria (2) automatically grants the criteria (3) $a_{3}>0$ and criteria (4) $a_{1}a_{2}a_{3}-a_{1}^{2}a_{4}-a_{3}^{2}>0$. Actually, as the satisfaction of criteria (2) in a classical model does not impose constraints on biomass (Eq.(28)-Eq.(29)), an important property in a classical model is that the stability of the system can be kept as long as the supply point [$R_{iron, supply}$, 0] locates within the region bounded by the reverse extensions of the consumption vectors.

However, in the iron-partition model, the self-creating dimension changes the sign of the coefficients $c_{11}$ and $c_{12}$. Under this situation, even when criteria (2) holds, criteria (3) and criteria (4) may still be violated. Due to the change in the sign of the coefficients $c_{11}$ and $c_{12}$, the occurrence of oscillations becomes possible, as well as exclusion and extinction, even when the supply point locates within the region bounded by the reverse extensions of the consumption vectors (Figure 3 and Figure S3).

### 4. Impact of parameters and the forms of equations on the results

#### 4.1. Discussion on the consumption rate, cost, and affinity of siderophores

Although we assumed in the main text that all public siderophores were recycled, modifying the model to allow the consumption of a portion of public siderophores does not affect the overall dynamics, including oscillation and stable coexistence. We also analyzed a modified version of Eq.(3) with a consumption ratio parameter $p$ ranging from 0 to 1 to indicate the proportion of consumption of the public siderophores:

$$\frac{dR_{\mathrm{sid}}}{dt}=\sum_{i=1}^{2} \frac{m_{i}}{r}\alpha_{i\text{, }\text{public}}\epsilon-p\sum_{i=1}^{2} \frac{m_{i}}{r}\cdot I\left( R_{\mathrm{iron}},R_{\mathrm{sid}},\vec{\alpha}_{i} \right)-dR_{\mathrm{sid}}$$

(37)

As shown by Figure S5, the higher the consumption ratio $p$, the easier for the partial cooperator to exclude or stably coexist with the cheater.

We also checked how the increases of production cost, i.e., the decreases of $\epsilon$, affect the systems’ dynamics. Intuitively, it becomes more difficult for the partial cooperator to coexist with the cheater (Figure S5).

We also examined the effect of affinity coefficients on the dynamics. Coexistence is possible over a wide range and becomes more likely in the oscillatory case as $K_{l}$ is smaller, which means the public siderophore has a much higher affinity for iron (Figure S6).

#### 4.2. Discussion on the forms of iron uptake fluxes

In our iron-partition model, we assumed the Monad form of iron uptake. There can be other forms of iron uptake:

Possibility (1): $J_{\sigma,\mathrm{private}}$ and $J_{\sigma, \mathrm{public}}$ can take the following mass-action forms instead of Monad form:

$$J_{\sigma,\mathrm{private}}=v_{m}\alpha_{\text{1,private }}\beta R_{\mathrm{iron}},$$

(38)

$$J_{\sigma, \mathrm{public}}=v_{l}R_{\mathrm{sid}}R_{\mathrm{iron}}.$$

(39)

Between strains adopting two strategies, at the possible intersection point, $R_{\mathrm{iron}}^{*}$ and $R_{\mathrm{sid}}^{*}$ can be solved as:

$$R_{\mathrm{sid}}^{*}=\frac{-\beta v_{m}\left( \alpha_{1, \mathrm{growth}}\alpha_{1,\mathrm{private}}{-\alpha}_{2, \mathrm{growth}}\alpha_{2, \mathrm{private}} \right)}{v_{l}\left( \alpha_{1, \mathrm{growth}}-\alpha_{2, \mathrm{growth}} \right)}.$$

(40)

$$R_{\mathrm{iron}}^{*}=\frac{-d\left( \alpha_{1, \mathrm{growth}}-\alpha_{2, \mathrm{growth}} \right)}{\beta\gamma v_{m}\alpha_{1, \mathrm{growth}}\alpha_{2, \mathrm{growth}}\left( \alpha_{1,\mathrm{private}}-\alpha_{2, \mathrm{private}} \right)},$$

(41)

Under this highly simplified form of iron-uptake fluxes, $R_{\mathrm{sid}}^{*}$ is a constant, and $R_{\mathrm{iron}}^{*}$ is a linear function with respect to $d$. That is, as $d$ increases, the intersection point moves parallel to the $R_{\mathrm{iron}}$-axis in the chemical space. It is obvious that $\alpha_{1, \mathrm{growth}}<\alpha_{2, \mathrm{growth}}$ is also the necessary condition for the intersection point to exist.

Possibility (2): When $J_{\sigma, \mathrm{public}}$ takes the following sigmoidal form of siderophores-iron complex:

$$J_{\sigma,\mathrm{private}}=\frac{v_{m}\alpha_{\text{1,private }}\beta R_{\mathrm{iron}}}{K_{m}+R_{\mathrm{iron}}},$$

(42)

$$J_{\sigma, \mathrm{public}}=\frac{v_{l}R_{\mathrm{sid}}R_{\mathrm{iron}}}{K_{l}+R_{\mathrm{sid}}R_{\mathrm{iron}}},$$

(43)

Between strains adopting two strategies, relationships between the intersection point $R_{\mathrm{iron}}^{*}$ and $R_{\mathrm{sid}}^{*}$ follows:

$$R_{\mathrm{sid}}^{*}=\frac{-\beta K_{l}v_{m}\left( \alpha_{1, \text{growth}} \alpha_{1,\text{private}}-\alpha_{2, \text{growth}} \alpha_{2, \text{private}} \right)}{v_{l}K_{m}\left( \alpha_{1, \text{growth}}-\alpha_{2, \text{growth}} \right)+R_{\mathrm{iron}}^{*}\left( v_{l}\left( \alpha_{1, \text{growth}}-\alpha_{2, \text{growth}} \right)+\beta v_{m}\left( \alpha_{1, \text{growth}} \alpha_{1,\text{private}}-\alpha_{2, \text{growth}}\alpha_{2, \text{private}} \right) \right)}$$

(44)

Despite that the form of $R_{\mathrm{sid}}^{*}$ is relatively complex, from the setting $\alpha_{1, growth}-\alpha_{2, growth}<0$, for a meaningful intersection to exist, Eq.(15) also must be satisfied. The solutions for $R_{\mathrm{iron}}^{*}$ and $R_{\mathrm{sid}}^{*}$ can be solved as:

$$R_{\mathrm{sid}}^{*}=-\frac{K_{l}\left( \alpha_{1,\text{growth}}\alpha_{1,\text{private}}-\alpha_{2,\text{growth}}\alpha_{2,\text{private}} \right)\left( \alpha_{1,\text{growth}}\left( \beta\gamma v_{m}\alpha_{2,\text{growth}}\left( \alpha_{1,\text{private}}-\alpha_{2, \text{private}} \right)+d \right)-d\alpha_{2,\text{growth}} \right)}{K_{m}\left( \alpha_{1,\text{growth}}-\alpha_{2,\text{growth}} \right)\left( \alpha_{1,\text{growth}}\alpha_{1,\text{private}}\left( \gamma v_{l}\alpha_{2,\text{growth}}-d \right)+\alpha_{2,\text{growth}}\alpha_{2,\text{private}}\left( d-\gamma v_{l}\alpha_{1,\text{growth}} \right) \right)}$$

(45)

$$R_{\mathrm{iron}}^{*}=-\frac{dK_{m}\left( \alpha_{1,\mathrm{growth}}-\alpha_{2,\mathrm{growth}} \right)}{d\left( \alpha_{1,\mathrm{growth}}-\alpha_{2,\mathrm{growth}} \right)+\beta\gamma v_{m}\alpha_{1,\mathrm{growth}}\alpha_{2,\mathrm{growth}}\left( \alpha_{1,\mathrm{private}}-\alpha_{2,\mathrm{private}} \right)}$$

(46)

### 5. The method of constructing the invasion chain

In obtaining Figure 4B, the simulated evolutionary process of strategies towards “evolutionarily optimal” can be expressed in the following steps:

(1) Arbitrarily assign a strain $\sigma$ with strategy $\vec{\alpha}_{\sigma}$ as the initial strategy. This initial strategy is selected as (1,0,0) in Figure 4B. If this strain can survive stably in the given chemostat condition, then it creates a steady-state chemical environment.

(2) Scan the whole strategy space for the “maximizing strategy” that maximizes growth rate under the current steady-state chemical environment. Then, the strain adopting this maximizing strategy is added to the system to invade. If the added strain can survive, the invasion is considered “successful”. The new chemical environment formed after adding new strains is used for another round of invasion (in the case of oscillatory coexistence, the environmental value of the instability point of the limit cycle is chosen).

(3) A set of strategies are considered as “evolutionarily stable” when the steady-state environment created by strains adopting these strategies cannot be invaded by any other strategies, i.e., no maximizing strategies could be found outside of the current strategies.

The invasion chain is formed by repeating (2) for successive invasions, till an evolutionarily stable environment can be achieved.

### Supplementary Figures

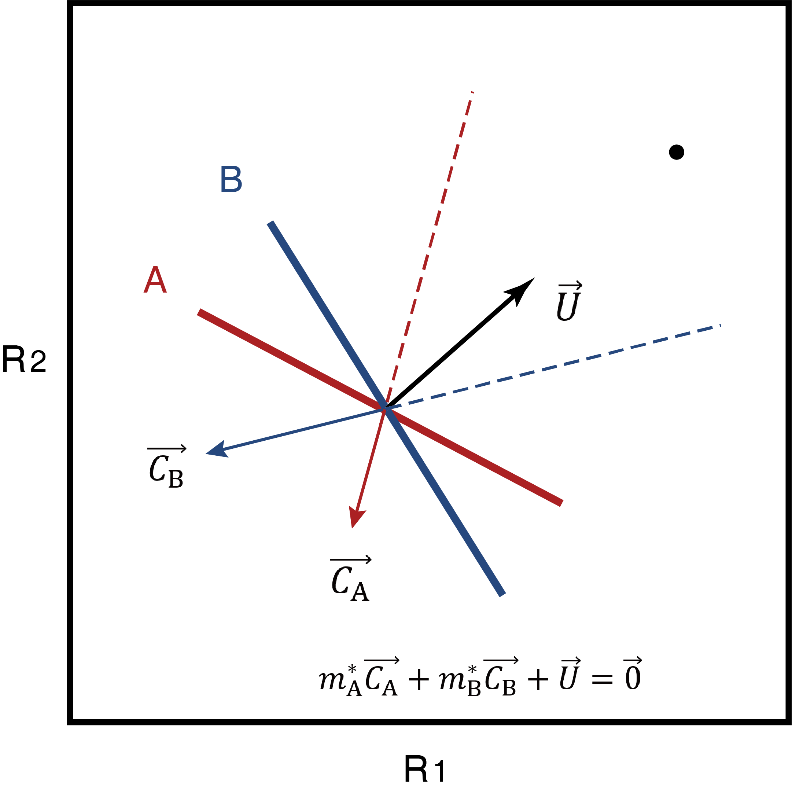

#### Figure S1. Diagram of the Tilman’s graphical tools on the classical consumer resource model

In the chemical space expanded by two resources $R_{1}$and $R_{2}$, The Tilman’s graphical tool in analyzing resource competition model contains:

(1) resource supply point $\left[ R_{1, supply},R_{2,supply} \right]$ (black dot) , which sets the maximal possible concentration of $R_{1}$ and $R_{2}$ that can occur at steady state;

(2) resource supply vector $\vec{U}$ (black arrow), which denotes the environmental supply rate of resources and $\vec{U}=d\left[ \begin{aligned} R_{1, supply}-R_{1}^{*} \\ R_{2,supply}-R_{2}^{*} \end{aligned} \right]$ at steady state;

(3) zero net growth isocline(ZNGI), or growth contour $\left\{ \left( R_{1},R_{2} \right) | \frac{{dm}_{i}}{dt}=0 \right\}$*,* for each strain (strain A: red line; strain B: blue line), which shows the contour where growth rate equals to death or dilution rate;

(4) consumption vector (red and blue arrows for that of strain A $\vec{C_{A}}$, and stain B $\vec{C_{B}}$, respectively), which indicates the total consumption rate of resources for the species. When two species reach a steady state, $m_{A}^{*}\vec{C_{A}}+m_{B}^{*}\vec{C_{B}}+\vec{U}=\vec{0}$. The consumption vectors and growth contours in this figure show that each of species A and species B consumes more of the one resource which more limits its own growth rate, leading to stable coexistence.

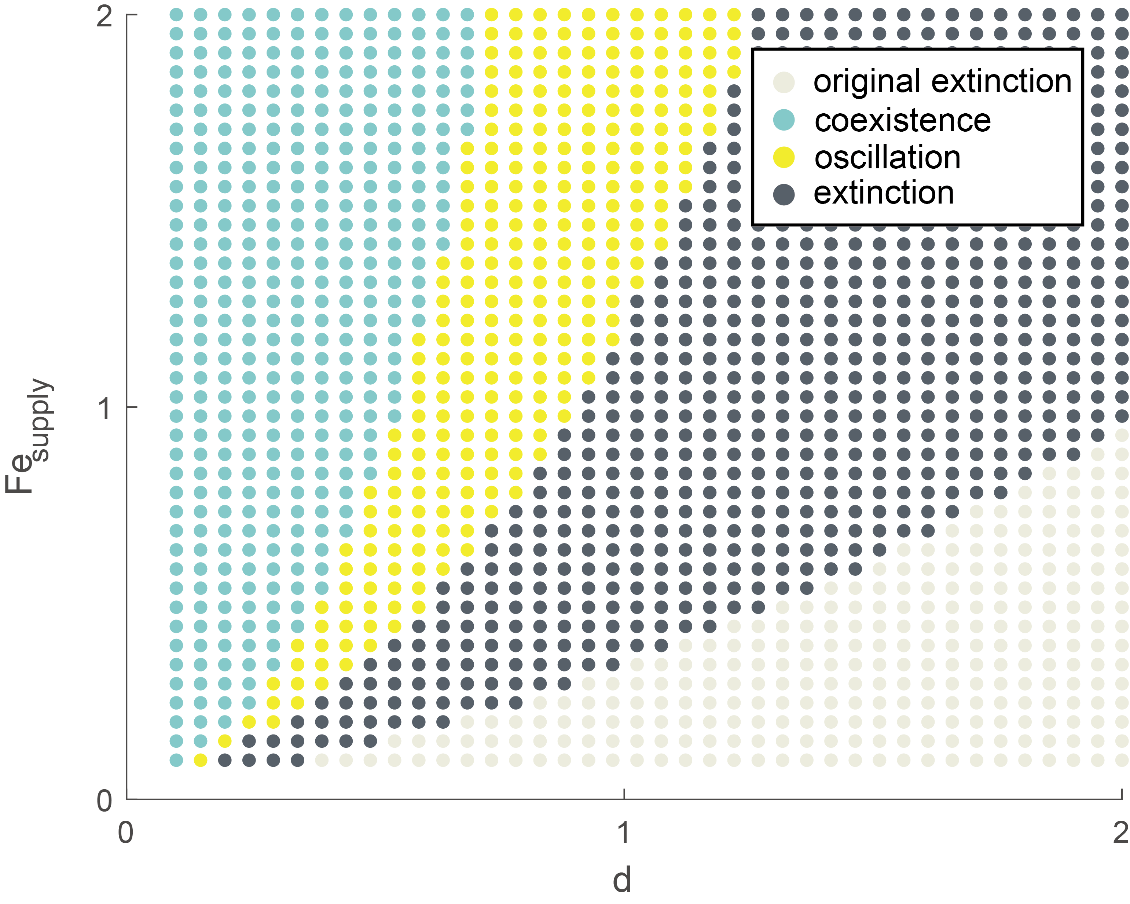

#### Figure S2. Cheaters accelerate the collapse of the system when invading partial cooperators.

Consequence of the pure cheater invading the partial cooperator $\vec{\alpha}=$(0.6,0.2,0.2) under various chemostat conditions. Light gray denotes chemostat conditions in which the partial cooperator cannot survive even on its own. Deep gray indicates regions where partial cooperators can exist on their own, yet the invasion of cheaters drives them to extinction. The yellow dots represent oscillatory dynamics, while the blue zone represents coexistence.

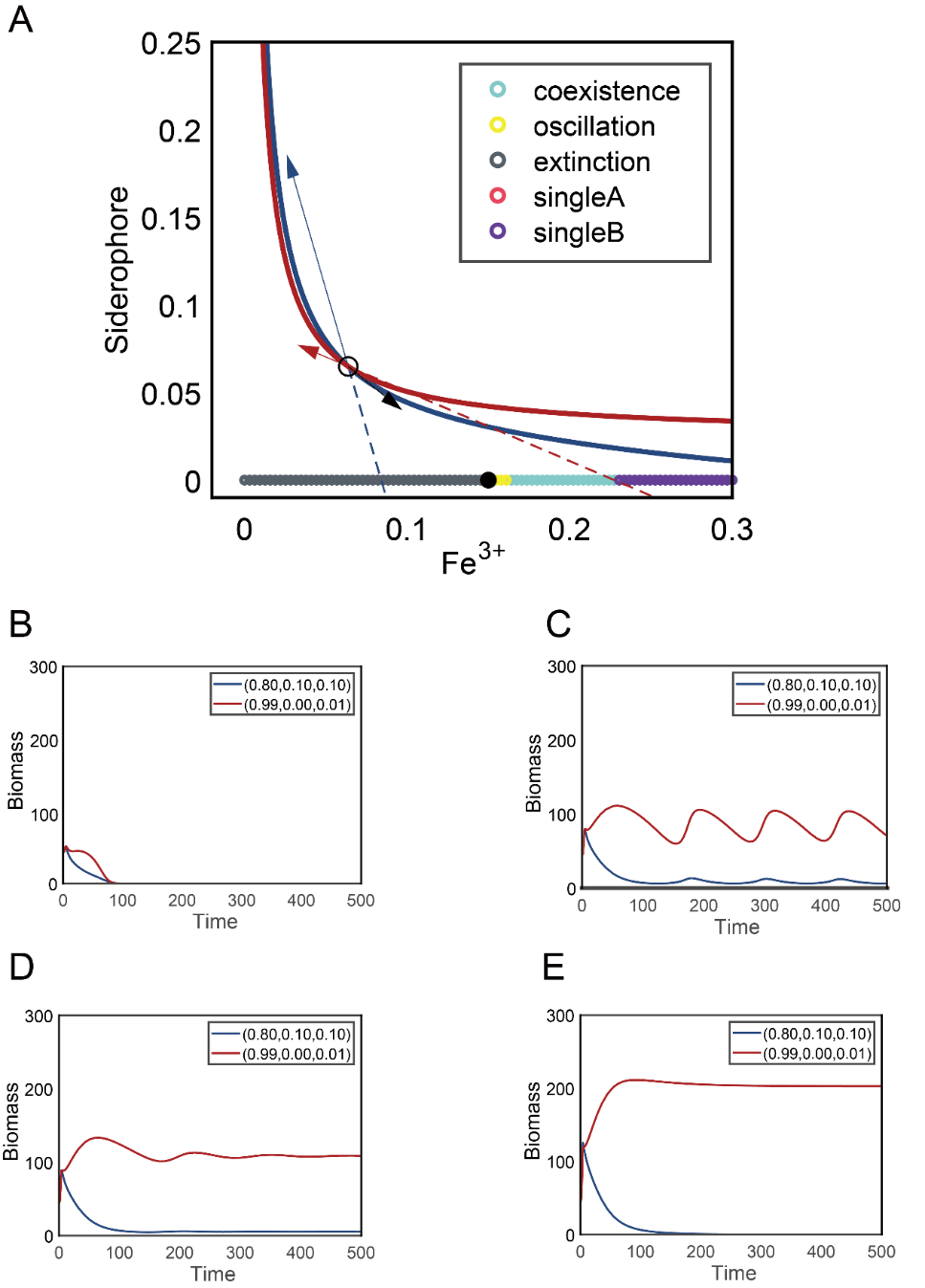

#### Figure S3. Competition between two cooperators under different Fe supply.

Similar as Figure 3 in the main text, but the system consists of two cooperators competing for iron. The strategies of the partial cooperator A is $\vec{\alpha}_{A}=$(0.8,0.1,0.1), and the nearly-pure cooperator B is $\vec{\alpha}_{B}=$(0.99,0,0.01).

(A) As the iron supply increases, the systems dynamics experiences extinction, oscillation, coexistence, and strain B excluding strain A, respectively. The interior of the reverse extension of the consumption vector covers the supply region where coexistence is possible, while the exterior is the region where exclusion occurs and the cooperator B survives alone.

(B-E) Time-courses of the competition dynamics between strain A and strain B under increasing iron supply, as shown in (A). (B): $R_{iron, supply}=0.1$; (C): $R_{iron, supply}=0.16$; (D): $R_{iron, supply}=0.18$; (E): $R_{iron, supply}=0.25$.

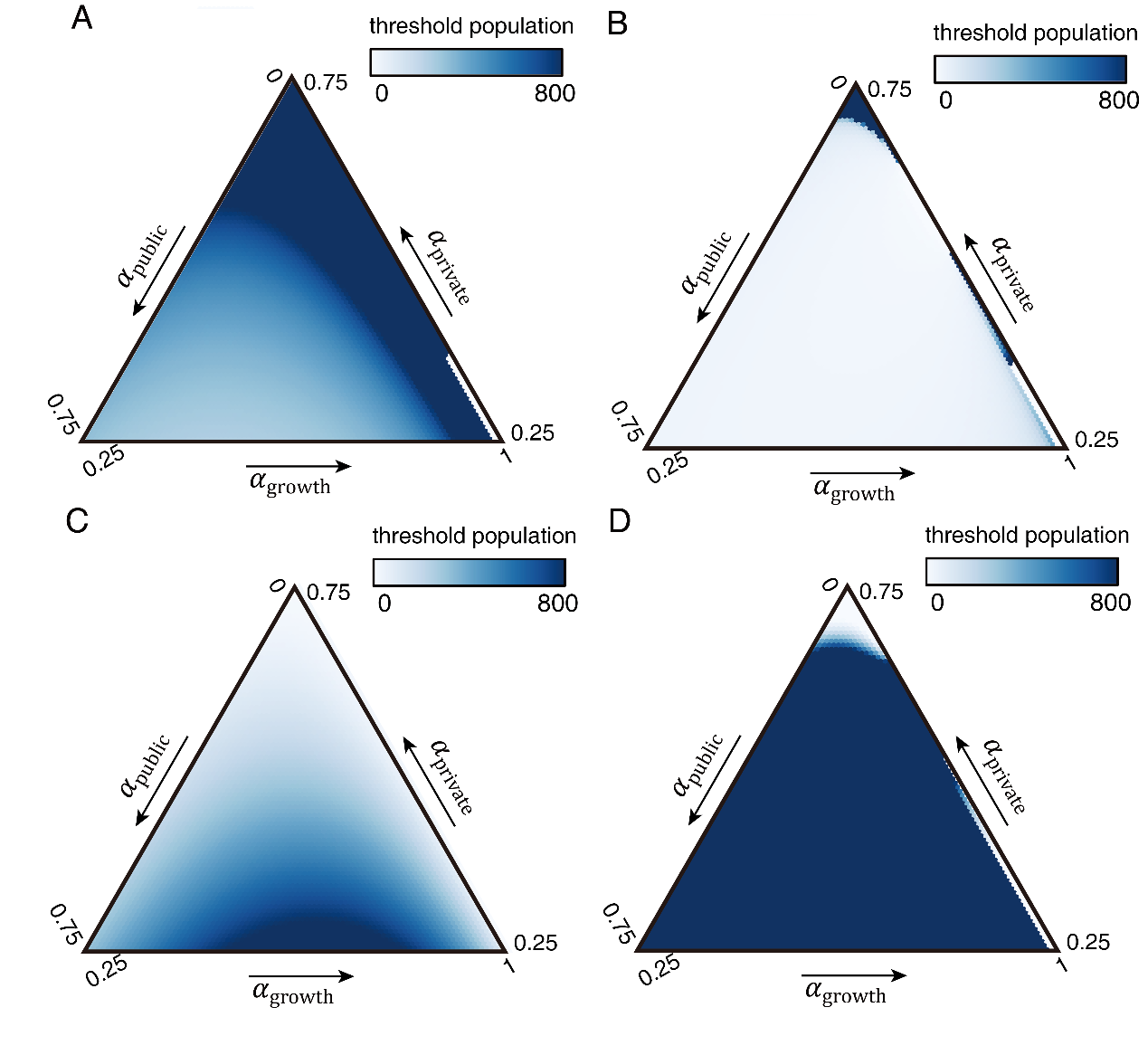

#### Figure S4. The invasion from another species with non-overlapping siderophores.

To assess a species' resistance to invasion by other species with different forms of siderophores, we modeled a second species with the same parameters but non-sharable siderophore. To assist the differentiation, we refer to the native species as species 1 and the invading species as species 2.

(A) The minimum population required for all strategies of species 2 to invade a native species 1 under the partial cooperator strategy ($\vec{\alpha}_{1}=$(0.6,0.2,0.2)).

(B) The minimum population required for all strategies of species 2 to invade a native species 1 under the self-seeker strategy ($\vec{\alpha}_{1}=$(0.6, 0.4, 0)).

(C) The minimum population required for a species 2 under the partial cooperator strategy(($\vec{\alpha}_{2}=$(0.6,0.2,0.2)) to invade all strategies of species 1.

(D) The minimum population required for a species 2 under the self-seeker strategy ($\vec{\alpha}_{2}=$(0.6, 0.4, 0)) to invade all strategies of species 1.

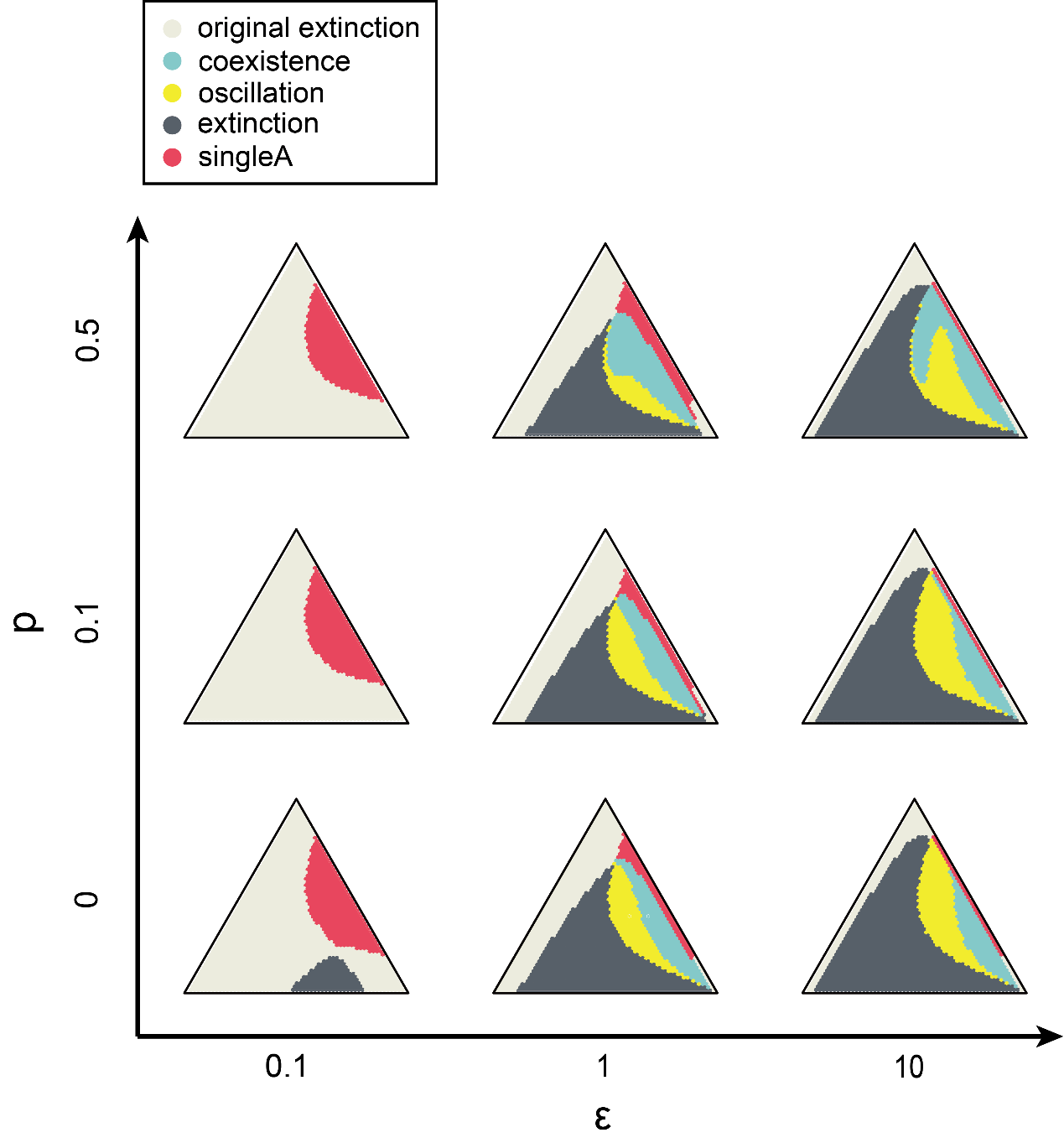

#### Figure S5. The impact of consumption, production, and recycle rate of public siderophores on strategies’ interaction consequences with the pure cheater.

To quantify the effects of siderophores’ production cost (represented by $\epsilon$) and recycle rate (represented by $p$) on its coexistence with the cheater, here we set up different combinations of $\epsilon$ and $p$ to reproduce Figure 4A. Along the x-axis, decreasing the production cost of public siderophores, i.e., larger $\epsilon$, helps to substantially enlarge the area of oscillation (yellow) and stable coexistence (blue) instead of exclusion (red). Along the y-axis, increased siderophore consumption, i.e., larger $p$, slightly increased the area of stable coexistence.

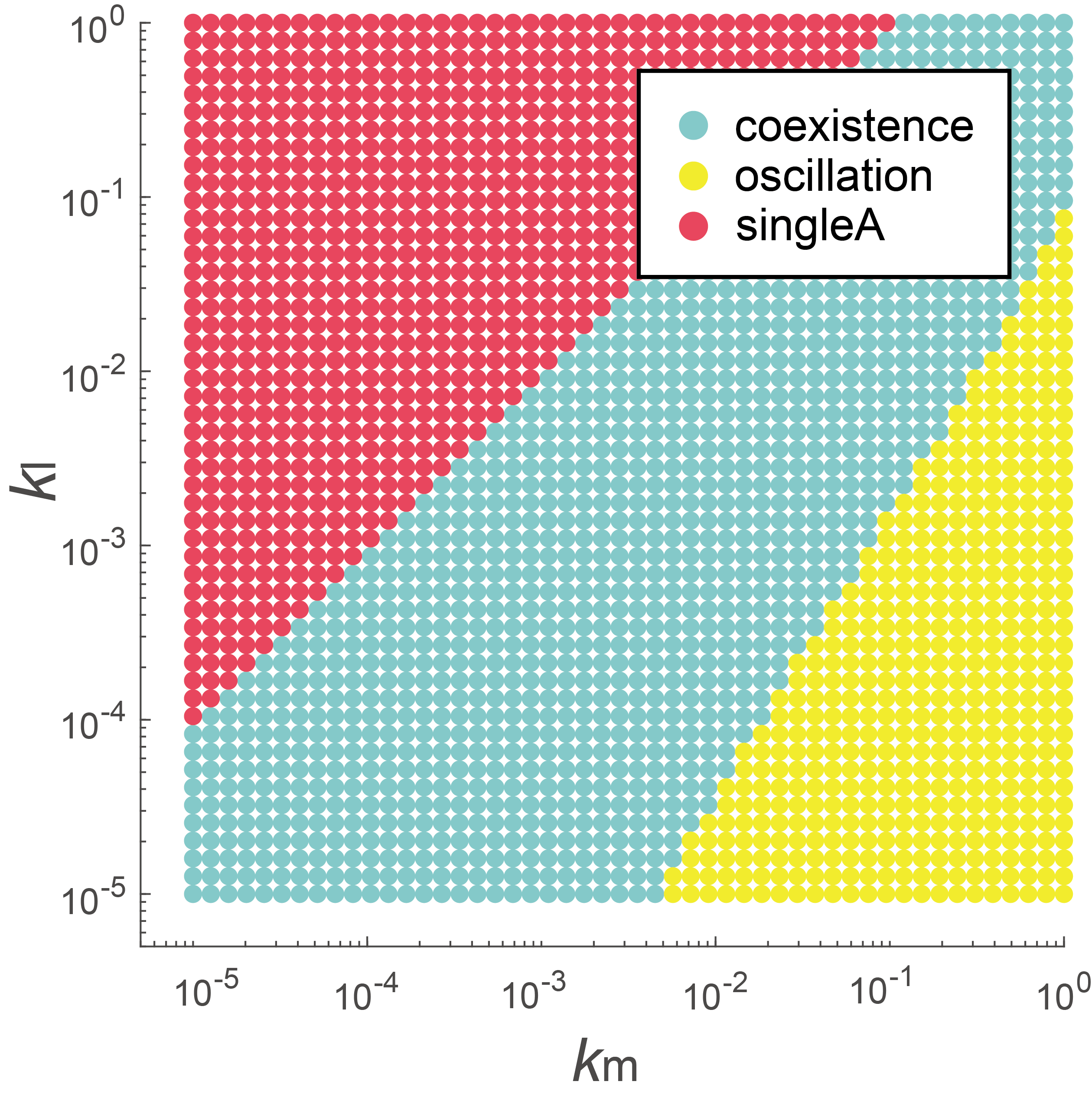

#### Figure S6. The impact of iron affinity of private and public siderophores on coexistence between the partial cooperator and the pure cheater.

$K_{m}$ and $K_{l}$ denote iron affinity of private and public siderophores, respectively. The consequence of the pure cheater invading a partial cooperator ($\vec{\alpha}_{A}=$(0.6,0.2,0.2), termed as “strain A”) under different levels of $K_{m}$ and $K_{l}$ were mapped to dot colors in the phase plane. Partial cooperators can stably coexist with pure cheaters in a wide range of parameter combinations, when $K_{m}$ balances with $K_{l}$(blue dots). When $K_{l}$ decreases and $K_{m}$ increases, i.e., public siderophores increase affinity for iron, the system is more likely to oscillate (yellow dot). When $K_{l}$ increases and $K_{m}$ decreases, the partial cooperator is more likely to exclude the cheater (red dots denoted as “single A”).

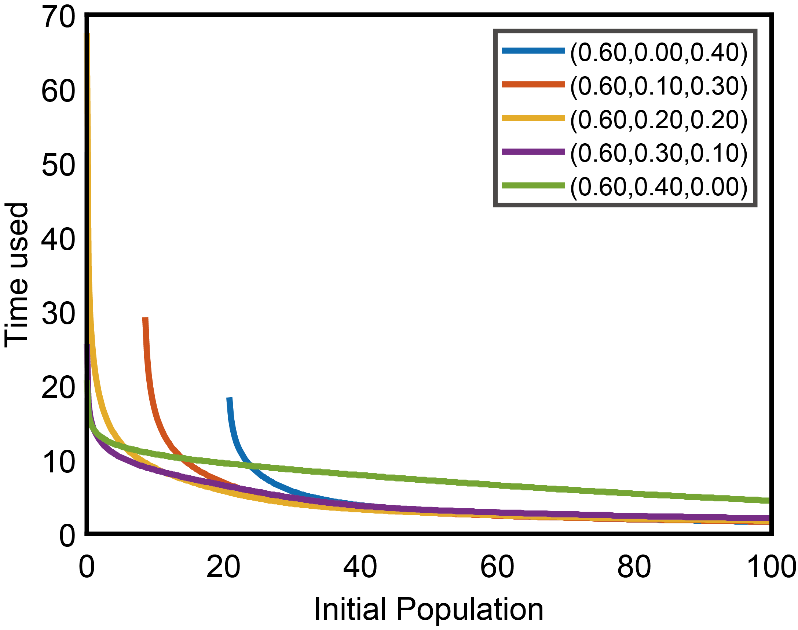

#### Figure S7. Speed of reaching a population's steady-state size for various strategies and initial populations.

In order to measure the speed of population establishment, we quantified the time used between introducing the initial population and reaching the steady-state population. Assuming constant $\alpha_{\mathrm{growth}}$, self-seekers and partial cooperators that allocate more resources to private siderophores can establish a population faster with a smaller initial population size, whereas strategies that allocate more resources to public siderophores can achieve the same or even faster results when the initial population size increases.

### Supplemental Table 3: Parameters used for different figures

| Term | Parameter values |
| --- | --- |
| Basic Parameter | $\beta=1$, $\gamma=10$, $\epsilon=1$, $K_{m}=1$, $K_{l}=0.1$, $v_{m}=1$, $v_{l}=1.$ |
| Figure 1 | $d=0.5$, $R_{iron, supply}=1.5$;  E: Pure cooperator: $\vec{\alpha}=\left( 0.6, 0, 0.4 \right)$;  F: Partial cooperator: $\vec{\alpha}=\left( 0.6, 0.2, 0.2 \right)$. |
| Figure 2 | A: Partial cooperator: $\vec{\alpha}=\left( 0.6, 0.2, 0.2 \right)$, $d=0.5$, $R_{iron, supply}=0.5$;  B: Partial cooperator: $\vec{\alpha}=\left( 0.6, 0.39, 0.01 \right)$, $d=0.5$, $R_{iron, supply}=0.5$;  D: Partial cooperator: $\vec{\alpha}=\left( 0.6, 0.2, 0.2 \right)$, $R_{iron, supply}=0.5$;  E: Partial cooperator: $\vec{\alpha}=\left( 0.6, 0.2, 0.2 \right)$, $d=0.5$. |
| Figure 3 | Partial cooperator: $\vec{\alpha}=\left( 0.6, 0.2, 0.2 \right)$, $d=0.5$  B: $R_{iron, supply}=0.3$; C: $R_{iron, supply}=0.4$; D: $R_{iron, supply}=0.6$ |
| Figure 4 | $d=0.5$, $R_{iron, supply}=0.5$,  Partial cooperator: $\vec{\alpha}=\left( 0.6, 0.2, 0.2 \right)$. B: start strain: $\vec{\alpha}=$ ($0.6, 0.2, 0.2$), middle strain: $\vec{\alpha}=\left( 1, 0, 0 \right)$, end strain: $\vec{\alpha}=\left( 0.65, 0.35, 0 \right)$ |
| S2 | Partial cooperator: $\vec{\alpha}=\left( 0.6, 0.2, 0.2 \right)$, $\epsilon=1$ |
| S3 | Partial cooperator: $\vec{\alpha}=\left( 0.8, 0.1, 0.1 \right)$,  Pure cooperator $\vec{\alpha}=\left( 0.99, 0, 0.01 \right)$*.*  B: $R_{iron, supply}=0.1$; C: $R_{iron, supply}=0.16$; D: $R_{iron, supply}=0.18$; E: $R_{iron, supply}=0.25$ |
| S4 | $d=0.5$, $R_{iron, supply}=0.5$ |
| S5 | $d=0.5$, $R_{iron, supply}=0.5$ |
| S6 | $d=0.5$, $R_{iron, supply}=1$  Partial cooperator: $\vec{\alpha}=\left( 0.6, 0.2, 0.2 \right)$ |
| S7 | $d=0.5$, $R_{iron, supply}=1$ |
